# Seasonal patterns of exometabolites depend on microbial functions in the oligotrophic ocean

**DOI:** 10.1101/2024.03.05.583599

**Authors:** Erin L. McParland, Fabian Wittmers, Luis M. Bolaños, Craig A. Carlson, Ruth Curry, Stephen J. Giovannoni, Michelle Michelsen, Rachel J. Parsons, Melissa C. Kido Soule, Gretchen J. Swarr, Ben Temperton, Kevin Vergin, Alexandra Z. Worden, Krista Longnecker, Elizabeth B. Kujawinski

## Abstract

Predictions of how the biogeochemical reservoir of marine dissolved organic matter (DOM) will respond to future ocean changes require an improved understanding of the thousands of individual microbe-molecule interactions which regulate the transformation and fate of DOM. Bulk characterizations of organic matter can mask this complex network of interactions comprised of rich chemical and taxonomic diversity. Here, we present a three-year, depth-resolved time-series of the seasonal dynamics of the exometabolome and the bacterioplankton community at the Bermuda Atlantic Time-series Study (BATS) site. We find both time-series to be highly structured and compositionally distinct across sampling depths. Putative exometabolite identifications (gonyol, glucose 6-sulfate, succinate, and trehalose) indicate that at least a portion of the exometabolome contains rapidly remineralized, labile molecules. We hypothesize that apparent seasonal accumulation of these labile molecules could result from environmental conditions that alter community composition on a seasonal timescale and thus shift the relative proportions of microbial functions that produce and consume the substrates. Critically, we found the composition of seasonal DOM features was more stable interannually than the microbial community structure. By estimating redundancy of metabolic functions responsible for cycling these molecules in BATS metagenomes, we propose a paradigm whereby core microbial metabolisms, either those utilized by all or by a subset of marine microbes, are better predictors of DOM composition than microbial taxonomies. The molecular-level characterization of DOM achieved herein highlights the metabolic imprint of microbial activity in DOM composition and greatly enhances our understanding of the dynamics regulating Earth’s largest reservoir of organic carbon.

**Significance statement:** Marine dissolved organic matter (DOM) is a major carbon reservoir that acts as a critical control on Earth’s climate. DOM dynamics are largely regulated by a complex web of microbial interactions, but the mechanisms underpinning these processes are not well understood. In a three-year time-series, we found thousands of DOM molecules and microbial taxa exhibited seasonal patterns. Critically, the identity of the microbes was more variable between years than the composition of the DOM molecules. We suggest that shared metabolisms encoded by genes that conduct core microbial functions are responsible for the more stable composition of DOM. This work links DOM molecules with microbial biodiversity, and presents testable predictors of DOM composition in our changing oceans.

## Introduction

In the global ocean, thousands of chemically diverse organic molecules are cycled by a rich and diverse microbial community. These interactions regulate the flux and storage of carbon in the biogeochemical cycling of marine dissolved organic matter (DOM) and thus exert critical controls on Earth’s climate (1). Both the chemical nature of DOM molecules (e.g., aromaticity, heteroatom content, size) and environmental conditions (e.g., community composition, microbial interactions, nutrient dynamics, temperature) have been proposed to control DOM flux (2–5). Disentangling the contributions of the different controlling mechanisms is key for predicting changes to DOM carbon flux in future oceans. To date, these mechanisms have most often been described by a framework of reactivity that condenses the thousands of DOM molecules into well-defined sub-pools. Labile and semi-labile DOM represent the most rapidly cycled substrates that sustain the microbial loop and result in carbon remineralization, whereas refractory DOM represents molecules that evade microbial degradation and sequester carbon in the oceans for thousands of years (1). However, critical dynamics of DOM are masked by bulk quantification (e.g., 6, 7).

Hundreds of thousands of individual microbe-molecule interactions leave a metabolic imprint on the standing stocks of DOM composition. These interactions form the fabric of the microbial web, supporting the relationships required to fix, exchange, metabolize, and ultimately remineralize carbon within the marine microbiome (8, 9). Thus, marine microbes act as the source and sink mechanisms of labile DOM flux. However, the cryptic nature of the DOM-microbe network inhibits our ability to predict when DOM composition will force a change in the microbial community or when microbial activity will alter the composition of DOM (10–12). This is further complicated by the vast diversity of DOM molecules and microbial taxa, but also their redundancies, where many different microbes can produce and consume the same DOM molecule. Nevertheless, key biogeochemical functions performed by marine microbes depend on the exchange of DOM molecules, and thus microbial taxonomy and metabolisms should be, at least partially, predictive of DOM molecular composition.

Previous culture experiments, as well as biogeochemical models, suggest that in similar environmental settings, the metabolic functions of microbial communities are more predictable than taxonomic composition (13, 14). This work reflects the fundamental nature of the core gene sets that encode the common metabolic functions of a taxonomic group. Overlayed on top of this is species or strain diversity that is created by adaptive traits not easily discerned in metabolic functions. While our understanding of microbial community assembly advances, a quantitative understanding of the relationship between these microbial drivers with their resulting metabolic by-products, or metabolites, that are released as DOM is lagging (15–19). Very recent marine metabolomics studies suggest that a limited number of marine metabolites are conserved across phylogenies, while others are taxonomically-specific (20–22). Parameterizing the relationships between DOM molecules and microbial community functions requires experimental efforts that simultaneously probe both the marine microbial community and DOM molecules and is essential for better-informed predictions of future ocean carbon cycling (23, 24).

To address this core challenge within marine biogeochemistry, we analyzed dynamics across a three-year, depth-resolved time-series of both DOM molecules and free-living microbial prokaryotic bacterioplankton in the seasonally oligotrophic northwestern Sargasso Sea at the Bermuda Atlantic Time-series Study (BATS) site. We present an unprecedented perspective into the seasonal variability of these two critical components of microbial-DOM interactions by disentangling bulk DOM into individual DOM molecules with untargeted exometabolomics analysis. Seasonal environmental changes represent recurring disturbances that induce shifts in the taxonomy and function of microbial assemblages (25), and thus create a natural perturbation ideal for testing the reproducibility of the resulting transformations in DOM composition. We assessed the seasonality and variability of both the exometabolome and bacterioplankton time-series with wavelet analysis. We show that within the BATS microbial community, the interannual variability of bacterioplankton taxonomy is greater than that of molecular-level DOM composition. This suggests that microbial assemblage mechanisms are functionally redundant so that the resulting DOM biogeochemistry remains consistent. With a targeted investigation of surface BATS metagenomes, we find that there can be a wide range in the degree of functional redundancy for enzymes responsible for producing and consuming seasonal exometabolites. Our work suggests that the presence and composition of exometabolites in the oligotrophic ocean will be determined by the presence of core metabolisms rather than by the presence of specific microbial taxa.

## Results and Discussion

With more than three decades of sustained observations, the large-scale biogeochemical and physical fields of the water column at the BATS site are well-defined (26–29). The BATS site is seasonally oligotrophic with recurring annual patterns of temperature and mixing in the epipelagic and upper mesopelagic. In winter and early spring, the system experiences convective mixing as deep as 200 – 300 m, where inorganic nutrients entrained from depth trigger an annual spring phytoplankton bloom (28, 29). Following the mixing period, a quiescent and stratified period develops in late spring and persists into mid-autumn, in which the surface 100 m becomes highly oligotrophic. These physical dynamics also drive seasonal dynamics of dissolved organic carbon (DOC), which is used as a proxy for bulk DOM, in the top 300 m at BATS (30, 31). During the stratified season (typically May – October), DOM accumulates in the euphotic zone (0 - 120 m). A portion of the seasonally accumulated, residual DOM is redistributed throughout the mixed layer and exported to the upper mesopelagic by deep convective overturning during the mixing season (typically January – March). Following re-stratification, the exported DOC becomes trapped in the mesopelagic, where it is subsequently remineralized by the resident microbial community (30–32). This physical framework guided our time-series sampling to capture the major water column states for the epipelagic and upper mesopelagic zones during all four major seasons (summer stratified, fall transition, winter mixed, and spring transition) (Fig 1A).

**Fig 1:**
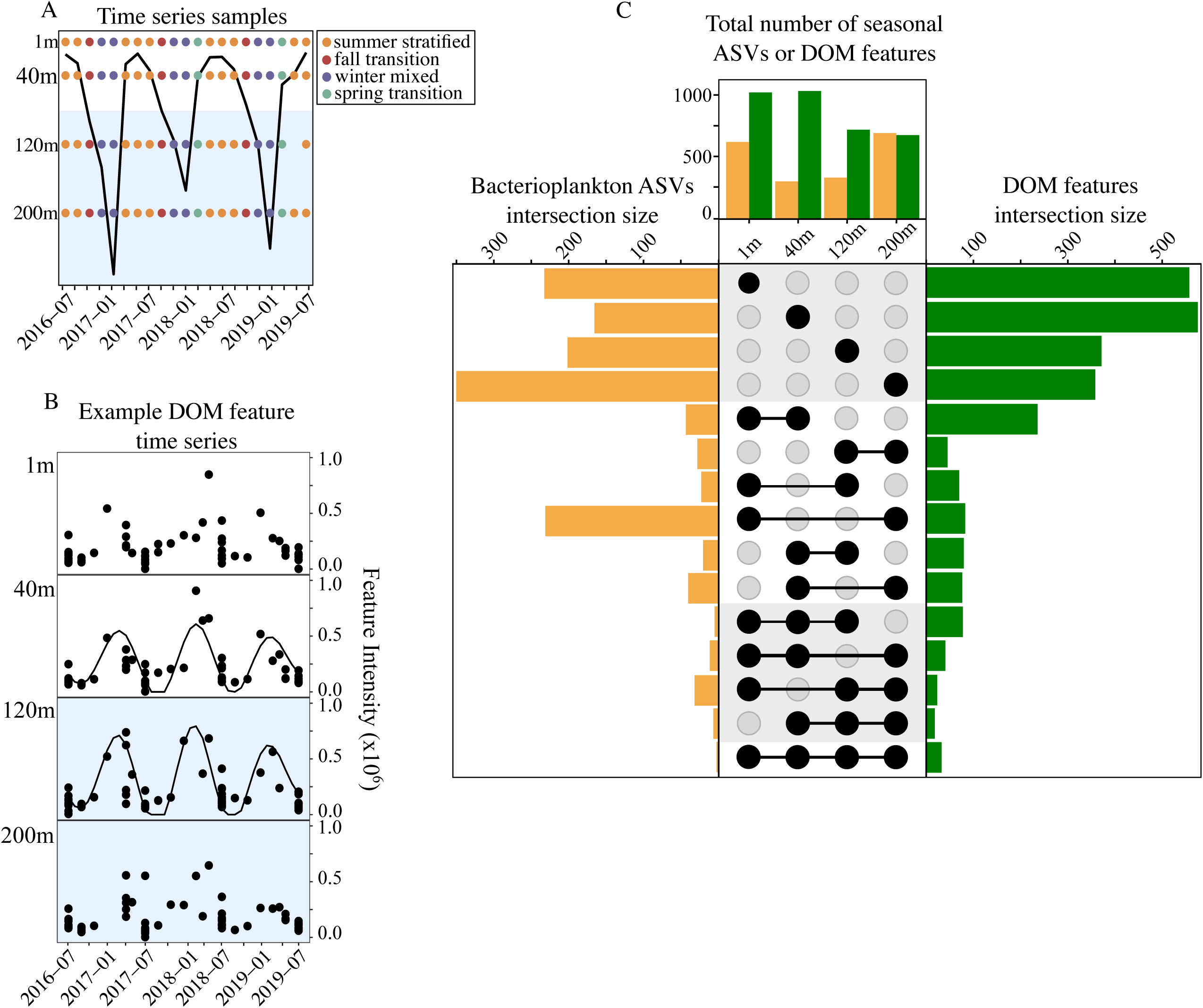
(A) Spatiotemporal coverage of samples collected in the three-year time-series for the BATS exometabolome and bacterioplankton community. Samples are colored by the physical framework seasons. The black line reflects mixed layer depth. The blue color indicates bottom of euphotic zone/transition to upper mesopelagic zone. (B) An example time-series of an unidentified DOM feature at all four sampling depths. The black dots reflect the feature’s intensity in all samples. Diel sampling efforts are reflected in the subset of days with multiple points displayed. At 40 m and 120 m, the grey line reflects the significant seasonal trend (12-month period) detected with wavelet analysis at these two depths. (C) Upset plot of depth structure in the seasonal bacterioplankton and exometabolome. The top bar plot reflects the total number of seasonal bacterioplankton ASVs (yellow) and DOM features (green) at each depth. Side panels reflect connectivity as defined by the inner legend. In the middle legend, black circles indicate the depth(s) at which seasonality was detected and the shading groups the different trends by the number of depths with connectivity. Rows 1-4 represent ASVs and DOM features that were seasonal at only one sampling depth, rows 5-9 represent seasonality at two sampling depths, rows 10-14 represent seasonality at three sampling depths, and row 15 represents seasonality at all four sampling depths. The bars reflect the sum of ASVs (left) or DOM features (right) that meet these criteria.

### Wavelet analysis detects seasonality in the exometabolome and bacterioplankton time-series

With the physical framework in mind, we collected a depth-resolved time-series of both the untargeted exometabolome, which represents a global overview of all DOM features detectable by solid phase extraction and liquid chromatography coupled to mass spectrometry, and the microbiome, using bacterioplankton V1-V2 16S rRNA gene amplicon sequence variants (ASVs). Parallel samples of DOM features and ASVs were collected bi-monthly for three-years from July 2016 to July 2019 at the surface (1 m), in the mixed layer (40 m), the base of the euphotic zone (120 m), and the upper mesopelagic zone (200 m) (Fig 1A) so that each DOM feature or ASV has an associated time-series at every sampling depth (e.g. Fig 1B). We detected 6293 DOM features, each defined by a unique mass-to-charge ratio and retention time. These DOM features were pre-filtered for peak quality, blank contaminants, isotopes, and adducts, and thus represent, to the best of our ability, a unique set of molecules. We compared the DOM features with patterns of 3158 bacterioplankton, defined as ASVs across the time-series which were pre-filtered to require presence in more than 5% of all samples. Key seasonal taxonomic trends in bacterioplankton succession at the BATS site have been previously described (33). Here we analyzed the temporal dynamics of the bacterioplankton dataset to compare with those of the untargeted exometabolome.

Due to the inherent challenges of comparing different data types across a time-series, we classified the temporal dynamics of DOM features and bacterioplankton ASVs using wavelet analysis to decompose each time-series at every sampling depth into the frequency domain (34) (Fig S1). Unlike clustering and correlation networks which can be used to group unknown DOM features based on temporal or spatial patterns (35, 36), wavelet analysis allows us to extract temporal insights of DOM molecules and ASVs that are concealed by these techniques, including the dominant periods (e.g., seasonal, 12 months), and the timing of period peaks (e.g., winter or summer). In addition, unlike some other time-series approaches, wavelet analysis can extract localized temporal information. This means that a period of interest does not have to occur globally across the entire time-series in order to be detected (37), and thus wavelet analysis can be valuable for detecting interannual differences in patterns of plankton communities (38). Here we compare the time-series of the untargeted exometabolome and bacterioplankton microbiome using wavelet analysis to analyze the spatial and temporal dynamics resulting from this complex network of microbe-DOM interactions.

The dominant period of a DOM feature or bacterioplankton ASV within the time-series was assigned based on the highest median power, an estimate of best fit, across all calculated periods (2-12 months). The best fit was required to be significantly different from a null hypothesis test of ‘no periodicity’ (median p-value ≤ 0.01) (Fig S2). We also required all time-series classified as having significant wavelets to have a relative standard deviation that represents a threshold for which environmental variability should be greater than analytical variability (> 25%) (39). Significant wavelets were found for 74% of DOM features (n = 4679) and 67% of ASVs (n = 2102) across the four sampling depths. The median powers ranged from 0.32 to 1.4, and the dominant periods ranged from 5 to 12 months (Fig S2). Almost all significant wavelets exhibited periods greater than 6 months, indicating that DOM features and bacterioplankton with shorter frequency periods were more stochastic and too similar to random white noise to be significant (Fig S2). Higher-resolution sampling and a longer time-series would be required to detect significant patterns with shorter frequencies. At almost every sampling depth, a majority of DOM features and ASVs exhibited a dominant period of 12 months, underscoring the important influence of seasonal environmental conditions (Fig S3).

### DOM features are differentiated by depth and season

A seasonal period of 12 months emerged as a dominant period across our time-series of the thousands of unknown DOM features that comprise bulk DOM, and the thousands of bacterioplankton ASVs that are, at least in part, responsible for the cycling of these molecules. A total of 2611 unique DOM features (41% of all) exhibited seasonality at one or more sampling depths (Fig 1C). Approximately two-fold more DOM features exhibited seasonality at 1 m and 40 m compared to DOM features at 120 m and 200 m (n = 1098, 1127, 700, and 665 seasonal DOM features, respectively). A total of 1385 unique ASVs (44% of all) exhibited seasonality at one or more sampling depths, and the greatest number of seasonal bacterioplankton was found at 200 m (n = 576, 291, 313, and 698 seasonal bacterioplankton at 1, 40, 120, and 200 m, respectively) (Fig 1C). To predict the season in which a seasonal DOM feature or bacterioplankton ASV time-series reached a maximum, wavelets were reconstructed with a period of 12 months and the season was assigned based on the month in which the maximum occurred (Fig S1C). Most seasonal DOM features (∼33-49%) and seasonal ASVs (83-95%) peaked in the summer stratified season at every sampling depth (Fig S4). The stratified periods encompass a large portion of the annual physical regime at BATS (Fig 1A), increasing the chances that our sampling would capture DOM features that exhibited a maximum during this period of elevated bulk DOM. However, we also found seasonal DOM features and bacterioplankton ASVs at each sampling depth in the time-series that peaked in the other seasons (fall transition, winter mixed, and spring transition) (Fig S4). The fewest seasonal DOM features peaked in the spring, though this is likely because more frequent sampling is required to capture the short-lived spring transition (Fig 1A). Bulk DOC exhibits a consistent seasonal cycle at BATS (4, 30). The exometabolome at BATS demonstrates that molecular patterns can reflect this bulk signal, as well as independent mechanisms. For example, a large majority of seasonal DOM features at 1 m (Fig S4) peaked during the summer stratified season, but an almost equal number of DOM features at this sampling depth exhibited peaks during the winter mixed season (e.g., Fig 1B). These peaks in winter correspond with the lowest DOC concentrations and thus represent new seasonal molecular signatures not detected by bulk methods.

DOM export can contribute significantly to carbon export in the subtropical oligotrophic ocean where deep convective mixing or subduction occurs (32, 40). For this reason, the connectivity of DOM features that exhibited seasonality at more than one sampling depth was of interest, particularly features that exhibited seasonality in both the surface (1 m or 40 m) and the deeper (120 m or 200 m) sampling depths as these DOM features could comprise a portion of DOC export. For example, the DOM feature in Figure 1B exhibited significant seasonality at both 40 m and 120 m. Although a small portion of DOM features exhibited seasonality in both the surface and deeper samples, the composition of the overall exometabolome was strongly vertically stratified and the transfer of seasonal DOM features between depths was limited (Fig 1C). The majority of the seasonal DOM features (n = 1840) exhibited seasonality at only one sampling depth (n = 540, 577, 366, and 357 seasonal DOM features were unique to the sampling depths of 1, 40, 120, and 200 m, respectively). A smaller subset of seasonal DOM features exhibited connectivity across depths, where the same DOM feature exhibited seasonality at two (n = 593), three (n = 148), or all four (n = 30) sampling depths. The greatest connectivity was found in DOM features exhibiting seasonality at both 1 m and 40 m (n = 239). Connectivity between the surface and deeper depths was minimal. Similar to the exometabolome, 85% of seasonal bacterioplankton ASVs (n = 862) exhibited seasonality in distinct depth zones (Fig 1C). A small subset of seasonal ASVs exhibited seasonality at two (n = 144) or three (n = 6) major depth zones. In contrast to DOM features, a subset of bacterioplankton (n = 231) peaked in the stratified season at 1 m and 200 m, indicative of some taxonomic connectivity in the surface and the upper mesopelagic bacterioplankton. Although depth is known to structure the ocean’s microbiome diversity (41), the influence on the resulting exometabolome’s spatiotemporal dynamics was previously unknown. The seasonal patterns of DOM features are highly stratified in the top 200 m and predominantly endemic to specific sampling depths.

Our individual time-series of thousands of DOM features detected temporal dynamics that complement trends in bulk DOC variability, but also independent dynamics that were unique to the exometabolome. The limited connectivity of seasonal DOM features was unexpected, given previous work, which finds that solid phase extraction retains the molecules that comprise exported DOM at BATS (36, 42). It is possible that a portion of these patterns result from depth-specific mechanisms, such as seasonal changes in zooplankton grazing or viral lysis (43, 44). This may also in part be explained by our Eulerian sampling approach and the interannual variability of the extent and duration of stratification and deep convection in our three-year time-series. The mixed layer extended deeper than 200 m in April 2017 and March 2019, but only to 174 m in March 2018 (Fig 1A). The different hydrographic conditions could have altered seasonal DOM compositions inconsistently between years at 200 m, which also happens to be the only sampling depth where wavelets of DOM features predominantly had a period of 8 months, rather than 12 months (Fig S3). The wavelet analysis is capable of detecting localized periods, but if these differences in winter mixing significantly disturb the periodicity of these DOM features, a different dominant period may be assigned. While the previously observed environmental controls of bulk exported DOM are seasonal, the rate of degradation of individual exported molecules is likely variable, and therefore a period of 12 months would not capture those dynamics. Thus, the seasonal exometabolome detected by wavelet analysis reflects a unique subset of the DOM reservoir.

Instead of recapitulating bulk DOM dynamics, the exometabolome at BATS captured new, additional patterns of seasonal behavior. A majority of the DOM features’ seasonal patterns were only observed within specific depth horizons. We posit that some of these individual molecules represent those that are cycled too quickly to be exported, resulting in their limited connectivity. Much of the bulk DOM that persists in the surface at BATS during summer stratification is considered to be semi-labile or semi-refractory as it is not accessible to the surface microbial communities, but can be degraded by genetically distinct microbial communities at depth after physical export by convective winter mixing (4, 45). While this mechanism is apparent in bulk DOC concentrations and characterized polymers, i.e., total hydrolysable amino acids (31), only a small portion of the seasonal DOM features were observed in both of our upper and lower sampling depths (Fig 1C). The seasonal peaks of a DOM feature in the exometabolome reflecting these bulk DOM patterns would be expected to differ in timing. The DOM feature would peak in the surface during summer stratification and subsequently peak in the deep during winter convective mixing. However, of the few DOM features that exhibited connectivity, most reached maxima in the same stratified season at both depths (Fig S4), again indicating that these seasonal DOM features were not persistent features redistributed by convective mixing. We hypothesize that these DOM features could be introduced as by-products of metabolisms that utilize the same metabolite across the different sampling depths or alternatively, other, more rapid export mechanisms such as sinking particle solubilization or the vertical-migrating mesozooplankton shuttle (46, 47).

### Labile exometabolites are present in the seasonal exometabolome

Untargeted exometabolomics techniques provide an opportunity to highlight important, but previously unrecognized, DOM molecules not detected with targeted techniques. Although the identification of environmental metabolites is a notoriously challenging endeavor (48), we highlight four putatively identified seasonal exometabolites of interest, which were identified to the highest levels of confidence possible (Level 1 or Level 2) (49): gonyol, glucose 6-sulfate (or the isomer galactose 6-sulfate), trehalose, and succinate (Fig 2, Table S1). All four exometabolites are presumably labile molecules based on structure and potential availability for microbial metabolism, exhibited seasonality in the surface, and peaked in the summer stratified season. Based on structures and existing literature, these four small exometabolites are expected to be rapidly metabolized. For example, gonyol has been shown to be consumed in 24 hours by the Alphaproteobacteria *Ruegeria pomeroyi* (50), and succinate is a widely used metabolite that acts as a crucial intermediate in the tricarboxylic acid cycle (see Supplementary Text 3 for further discussion of potential microbial interactions). Thus, these exometabolites have the potential to play an important role in the ocean’s microbe-DOM network. In addition, based on previous work with structurally similar molecules, we assume that the four molecules have low extraction efficiencies (∼ <1%) when using solid phase extraction and therefore must be present at high concentrations to be observable in this time-series (51). The identification of these exometabolites highlights their importance, and future efforts using targeted extraction techniques (e.g., 42, 43) will further elucidate absolute concentrations to bolster the current paucity of *in situ* observations.

**Fig 2:**
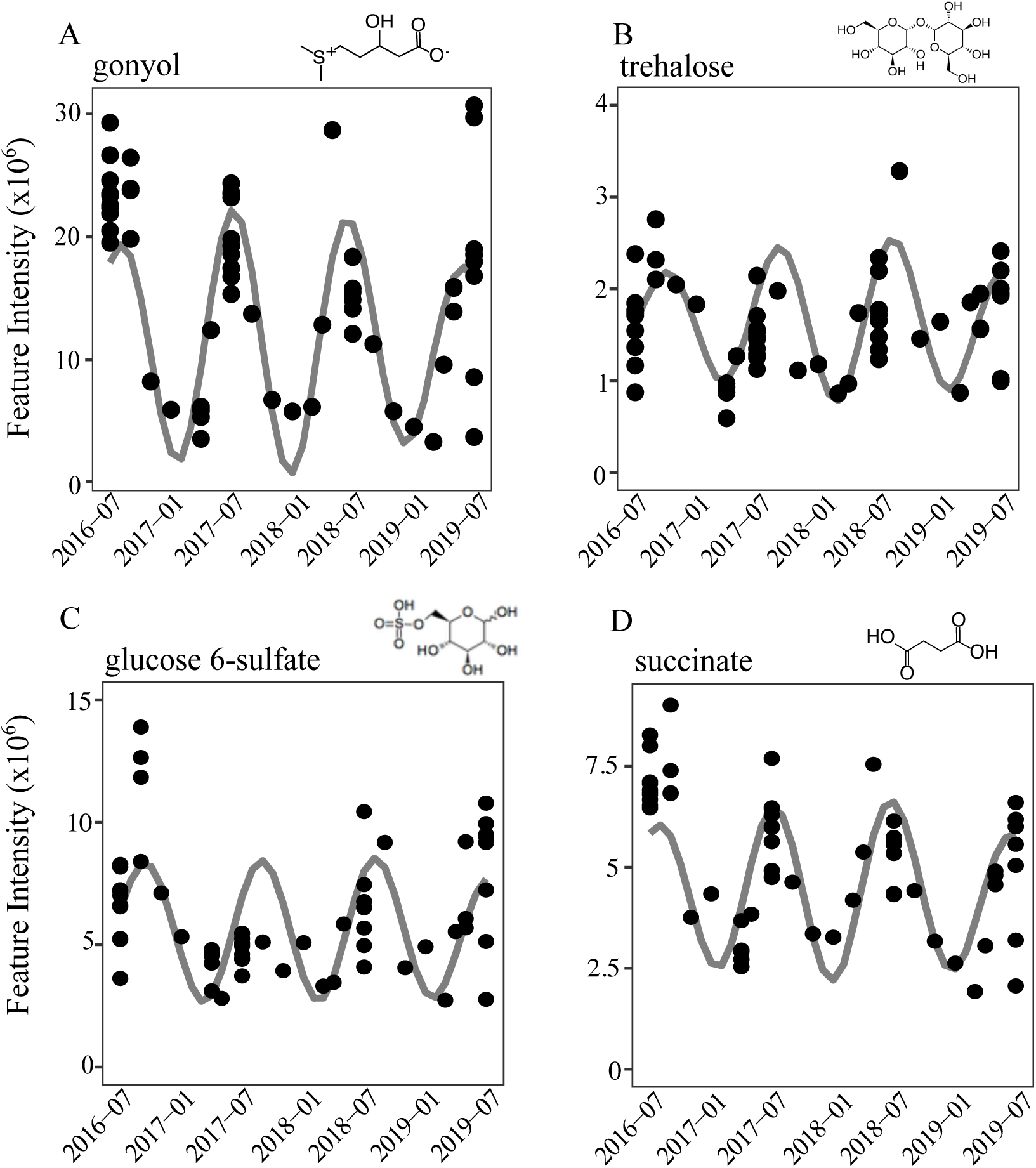
Seasonal patterns across the three-year time-series of four putatively identified exometabolites: gonyol, glucose 6-sulfate (or the isomer galactose 6-sulfate), trehalose, and succinate. All samples collected are presented (black circles), including July diel campaigns. The significant seasonal pattern (grey lines) was calculated with a reconstruction of the wavelet using a 12-month period. Feature intensity units are arbitrary (see Methods). The presented seasonal patterns are from sampling depth 1 m.

The composition of most of the seasonal exometabolome remains unknown. Solid phase extraction is known to select for the more recalcitrant-like properties of bulk DOM based on both the resulting composition of extracted DOM and its potential for microbial degradation (54). For example, the size and C:N ratio of DOM decreases after solid phase extraction, and the drawdown of bulk DOC concentrations are lower when DOM is provided to microbial communities after solid phase extraction as compared to fresh DOM (55–57). However, as described above, our time-series captured a unique subset of seasonal DOM that is vertically stratified and specific to sampling depths. By looking at individual molecules, we found that labile exometabolites do exist within the seasonal exometabolome, and quantitative targeted work of the same time-series found additional labile exometabolites (n = 14) that also exhibit seasonal behavior at BATS (58).

Certainly not all, and not even a majority, of these seasonal DOM features are expected to be labile, but based on this work, and previous studies (58–61), exometabolomes collected using solid-phase extraction contain a vast range of molecular compositions and individual components have the potential to be rapidly remineralized.

### Seasonal patterns of DOM features break away from bulk DOM time-series

The mechanisms controlling bulk DOM recycling have traditionally been defined by a spectrum of turnover times and reactivities, from the most labile and reactive DOM to the most recalcitrant and long-lived DOM (1). In the context of the bulk DOM framework, seasonal DOM features in this time-series would be classified as semi-labile or semi-refractory because of their apparent seasonal accumulation (62). However, our putative identifications indicate the presence of labile molecules within this seasonally-accumulating pool. This summer accumulation has been observed at BATS for other similarly small and easily metabolized molecules detected with targeted methods (58, 63), which can have rapid half-lives of ∼24 hours (64). Signatures of these individual DOM molecules are obfuscated within the µM resolution of bulk DOC methods. When tracking individual DOM molecules, it should be expected that these molecules can diverge from the bulk DOM framework and that observations of exometabolite patterns may result from independent mechanisms.

Vertical stratification of microbial taxa in the oligotrophic ocean presumably results from niche partitioning among microbial specialists along nutrient and energy gradients (65–67). Culture-based work has shown certain exometabolites are released uniquely by distinct phylogenetic groups and strains (22, 68). The exometabolome time-series at BATS provides *in situ* evidence that vertical differentiation of the microbial community also promotes stratification of metabolic by-products. Based on the timing of seasonal peaks and the depth differentiation of seasonal DOM features, the BATS exometabolome likely captured molecules with abiotic and biotic control mechanisms unique to each molecule. Conservative dilution in the surface and physical export below the euphotic zone may produce the observed patterns (30, 32). The persistence of these molecules may result from inherent recalcitrance or environmental conditions that do not support microbial communities capable of accessing the molecules. However, the unique seasonal patterns of DOM features that were independent of bulk DOM dynamics support the hypothesis that the exometabolome captured some of the rapid metabolic rate processes of the microbial communities, rather than just the production of persistent molecules. We propose that some of the observed patterns are the result of biotic controls due to rapid microbial turnover. The apparent seasonal accumulation of DOM features could emerge from seasonal changes in community composition and unequal shifts in the expression of production and consumption processes.

### Exometabolome composition is more stable interannually than microbial taxonomy

During the three-year time-series, we tested whether the observed temporal dynamics of the exometabolome and bacterioplankton community remain consistent between years. We assessed interannual variability using the wavelet’s median power (Fig S1), which reflects its fit and thus provides insight into the predictability of a given time-series (Fig 3). A high median power indicates the time-series fits well to the seasonal wavelet. A low median power can be driven by a poor fit and/or a signal that only appears in a portion of the three-year time-series. Across all sampling depths, the range of median powers of the seasonal DOM features and bacterioplankton were similar (0.3 - 1.3). However, the distributions of median powers were significantly different for DOM features than for bacterioplankton (Kolmogorov-Smirnov test D = 0.2, p < 2e-16) (Fig 3). Seasonal DOM features had an average median power of 0.6 ± 0.2 (mean ± std dev), which was 0.1 units greater than the average median power of bacterioplankton (0.5 ± 0.1), indicating that DOM features exhibit stronger recurring patterns between years at BATS. Additionally, the consecutive absences, or sparsity, in the two data types supports our observation that interannual variability was greater in seasonal ASVs than in seasonal DOM features. Across all seasonal ASVs, ∼30% (n = 597) were completely absent in at least one year of sampling in the time-series. In comparison, only a few seasonal DOM features (n = 8) contained consecutive zeros across one of the years sampled in the time-series.

**Fig 3:**
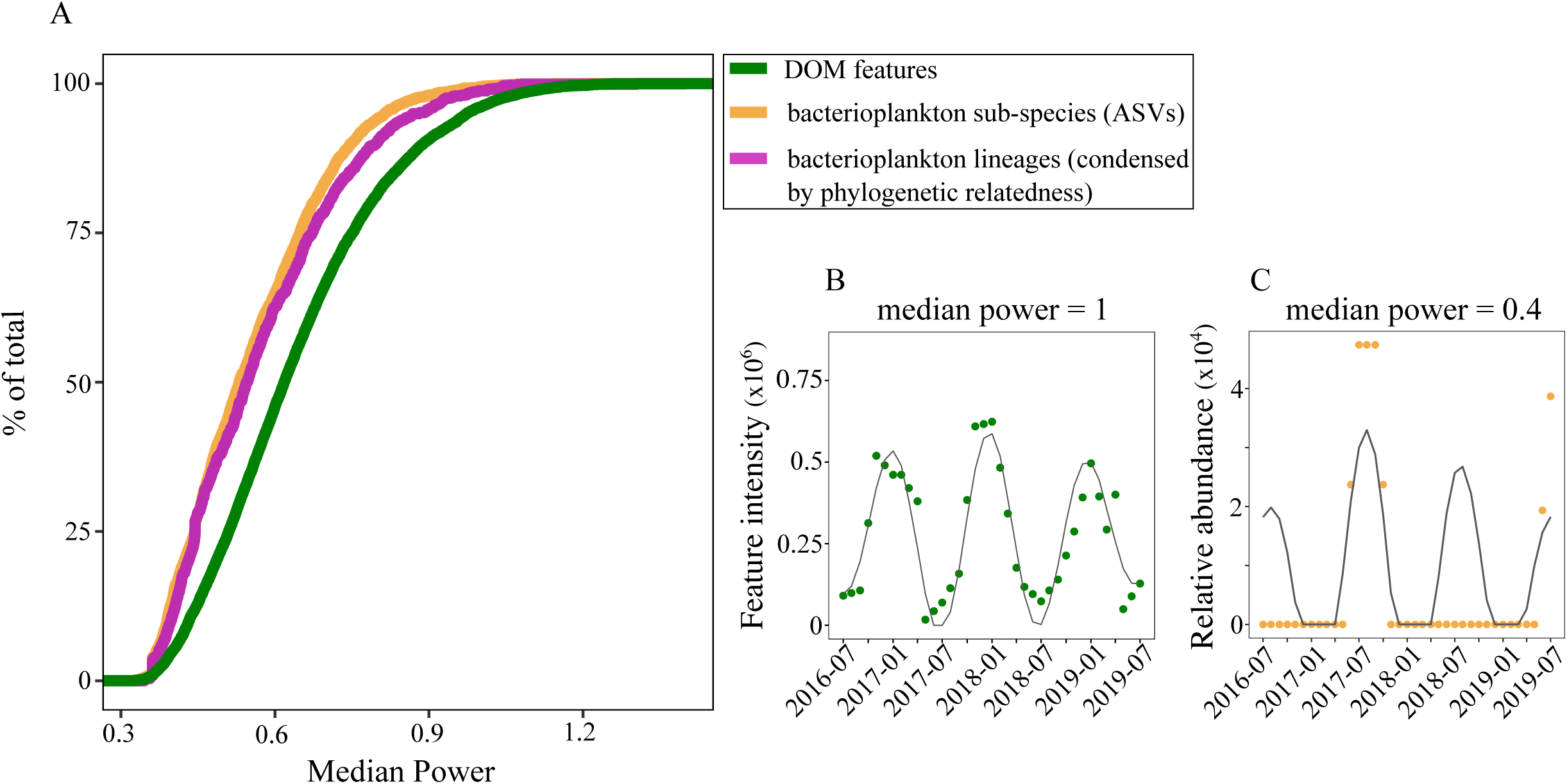
(A) Empirical cumulative distribution functions demonstrating the spread of median powers calculated for seasonal DOM features (green), seasonal bacterioplankton (defined as ASVs) (yellow), and seasonal bacterioplankton nodes condensed by phylogenetic-relatedness (pink). The insets demonstrate two examples of time-series with different median powers. (B) Example of DOM feature with a high median power, which reflects a seasonal wavelet fit that is predictable and exhibits the same pattern across the three-years. (C) Example of bacterioplankton ASV with lower median power, which is reflects a poor wavelet fit and absences across the three-years.

The magnitude of richness between the DOM features and bacterioplankton ASVs being compared are similar. However, we also compared the distributions of median powers with the time-series of ASVs condensed to node-resolved taxonomic resolution based on phylogenetic relationships to confirm that the interannual variability observed for the bacterioplankton was not inflated by an overrepresentation of rare taxa in ASV microdiversity (69, 70). Even after condensing the highly resolved ASVs, the same difference in the distribution of median powers was observed (Fig 3). This suggests that distinct microbial taxa, rather than just slightly different ASVs, are responsible for the interannual changes observed across the bacterioplankton time-series.

The composition of seasonal DOM features remained statistically stable across the three years despite changes in the bacterioplankton community (Fig 3). This indicates that some form of metabolic redundancy across the variable taxa promoted a stable state of equilibrium in DOM composition across the time-series (71). Other time-series studies have found that individual microbial taxa can vary between years (72, 73), but the resulting feedback on the exometabolome composition was previously unknown. Here we show that the controls of taxonomic variability in ASVs and their 16S rRNA gene differ from those of the exometabolome, making it difficult to utilize highly-resolved taxonomic information in predictions of DOM composition.

### Redundant metabolisms underpin the seasonal exometabolome

We hypothesized that recurring patterns of the same DOM features in the exometabolome result from a bacterioplankton community that exhibits taxonomic variability across years but convergent functionality of key metabolisms. Marine microbial communities are extremely diverse, but also share core groups of genes that confer the same functions (41). Core genes can be defined as a set of genes that are common to a single species based on different strains’ genomes or to an entire microbial community within a specific environment based on metagenome samples (41, 74, 75). This functional redundancy of specific enzymes or metabolic pathways is thought to create a buffering capacity of microbial ecosystem functions when communities change (76, 77). To test our hypothesis, we conducted targeted analyses of historical metagenomes and quantified the redundancy of metabolic reactions that involve trehalose or succinate as a product or reactant. These two exometabolites represent examples of DOM features that exhibited consistent interannual seasonal patterns in the surface exometabolome at BATS, but also their utilization is expected to differ significantly across the microbial community. Succinate is broadly used as part of the tricarboxylic acid (TCA) cycle, whereas trehalose is used more narrowly as a carbon substrate or for energy storage (78).

Genes that perform reactions involving the identified exometabolites were searched in metagenomes to assess the redundancy of succinate and trehalose metabolisms (Table S2). Thus, here we define functional redundancy as the degree to which a sequence is present across a metagenome. Genes were surveyed using functional orthologs (KOs) in 22 years (1997–2019) of all known publicly available surface ocean BATS metagenomes (n = 28 samples) (Table S3). These samples were not uniformly collected, but they capture all four seasons of the BATS physical framework (sample numbers from each season are n = 12 summer stratified, 5 fall transition, 9 winter mixed, and 2 spring transition) (Table S3). In cases where more than one KO exists for a given reaction (Table S2), we only present the most common KO (Fig 4).

**Fig 4:**
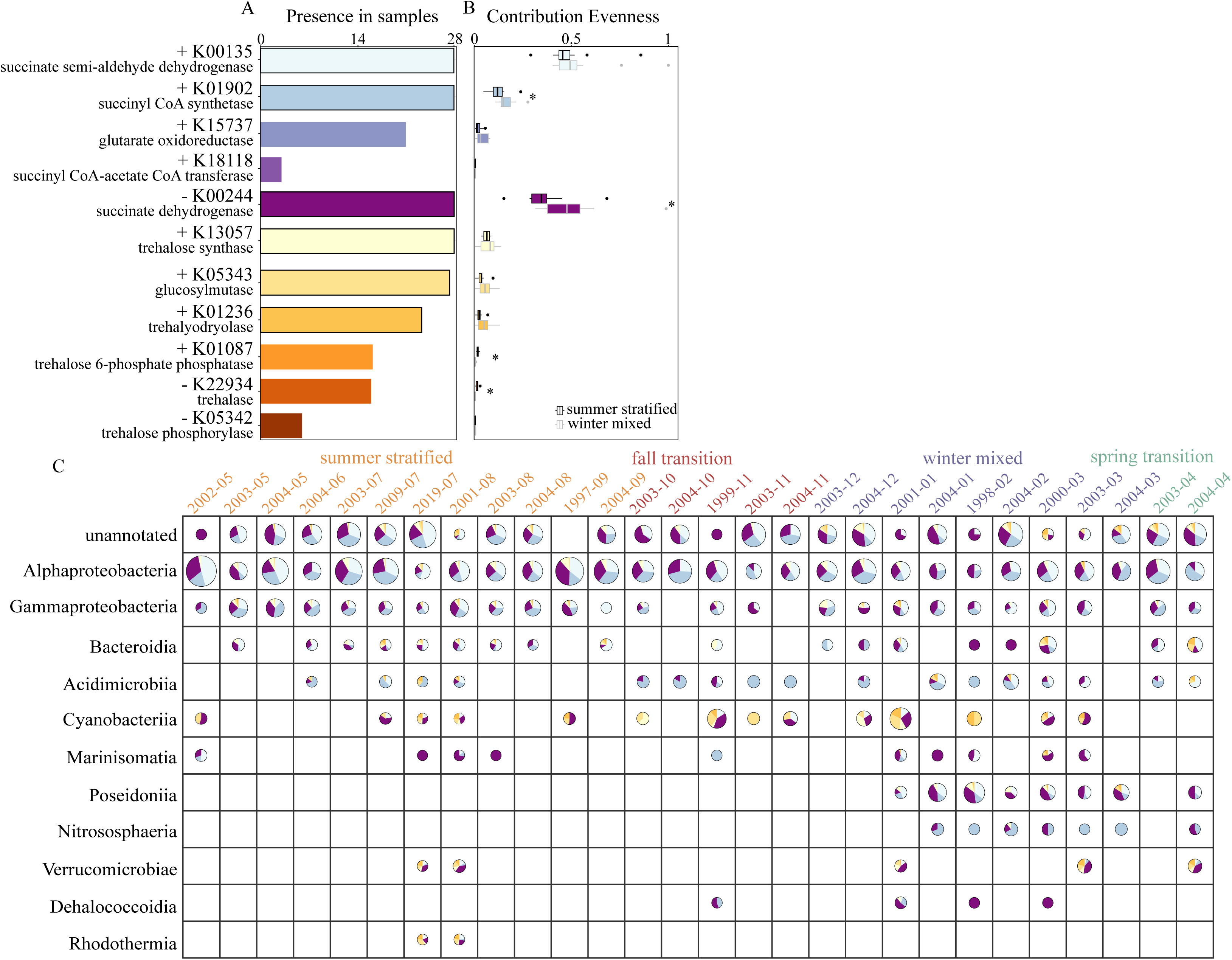
(A) Presence of genes as represented by metabolic functions (KOs) that utilize succinate (cool colors) or trehalose (warm colors) as a product or reactant in surface metagenomes at BATS (n = 28 samples total). + indicates exometabolite is product. – indicates exometabolite is reactant. The six KOs defined as being core genes are outlined in black. (B) The functional redundancy of the same genes as estimated with the metric of contribution evenness (CE) of each gene. Higher CE values reflect greater functional redundancy. The top boxplot (black outline) are CE values in metagenomes collected during the summer stratified season and the bottom boxplot (grey outline) collected during the winter mixed season. A star indicates CE was significantly different between seasons (Wilcoxon rank sum test, p ≤ 0.1). (C) The pie charts reflect the functional taxonomy of the same genes. The most commonly annotated (present in ≥ 2 samples) classes are presented (rows) in each of the metagenome samples (columns) as organized by season (summer stratified, fall transition, winter mixed, and spring transition). Each pie is divided by the relative contribution of the taxonomic group to each of the six core genes (K00135, K01902, K00244, K13057, K0543, K01236) based on total RPKM (reads per kilobase per million mapped reads) in a given sample. KO colors are the same as presented in (A). The pies are scaled based on the total relative contribution of each taxonomic group to the sample.

Most succinate genes (n = 5 reactions) and trehalose genes (n = 6 reactions) were present in a majority of the metagenome samples (Fig 4A). We found genes for both succinate (n = 3) and trehalose (n = 3) that were present in 100% and >85% of surface metagenome samples, respectively, and thus assume that these are core genes of the surface microbial community at BATS (Fig 4A). We estimated functional redundancy in the surface microbial communities with the metric of contribution evenness (CE) based on gene abundances (79) (Fig 4B). CE ranges from no redundancy (CE = 0), indicating only one community member in the sample harbors the gene of interest, to absolute redundancy (CE = 1), indicating all community members contribute equally to the presence of the gene of interest. As would be expected based on the increase in sequencing power in the last twenty years, sequencing depths varied by orders of magnitude across the different metagenomes. CE accounts for these differences by normalizing KO abundances to total species richness as estimated by the presence of universal single-copy marker genes, which are assumed to occur once in each genome (Fig S5). The maximum CE value for succinate-related genes was 1 for both K00135 and K00244, reflecting the ubiquity of the TCA cycle. In comparison, the maximum CE value for trehalose-related genes was 0.13 for K13057, reflecting the narrower potential for trehalose utilization in marine microbial communities. The median CE across all samples ranged from 0 to 0.5 for all succinate-related genes and from 0 to 0.1 for all trehalose-related genes (Fig 4B). CE of succinate-related genes were overall significantly greater than those of trehalose-related genes (Wilcoxon rank sum test p ≤ 0.01), indicating greater redundancy in succinate metabolism.

Functional taxonomy of these genes reflected the differentiation of the microbial community’s ability to utilize succinate or trehalose (Fig 4C). Core genes encoding for enzymes required to conduct the TCA cycle (K00135, K01902, and K00244) were most commonly annotated as Alphaproteobacteria, specifically *Pelagibacter,* and Gammaproteobacteria, specifically SAR86, which accounted for ≥ 67% and ≥ 15% of the sum of annotated RPKM for each core KO. As these are dominant groups in the surface ocean at BATS (80–82), it is not surprising that they dominate the taxonomy of genes required for the widely used TCA cycle. In contrast, the functional taxonomy of core genes encoding for enzymes that synthesize trehalose (K13057, K05343, and K01236) was more specific. The Alphaproteobacteria were also the most commonly annotated taxonomic contributors of the trehalose synthase K13057 (accounted for 64% of the sum of annotated RPKM), whereas the other two core trehalose genes (K01236 and K05343) were most commonly annotated as Cyanobacteria or Bacteroidia, which accounted for >31% and 15% of the sum of annotated RPKM for each KO. Although the extent of functional redundancy and the dominant functional taxonomy differ between succinate and trehalose metabolisms, both metabolisms are associated with multiple core genes in the surface microbial community and also very similar resulting patterns of exometabolites.

Between seasons, both functional redundancy and functional taxonomy (Fig 4) of production and consumption proteins exhibited shifts that could be attributed to the biological controls of the seasonal DOM features we observed (Fig 2). A subset of the analyzed succinate and trehalose genes exhibited no significant difference in CE between the summer stratified and winter mixed seasons (Fig 4B), but there was a general taxonomic shift in their major contributors based on its presence in samples (Fig 4C). For example, CE of the most redundant succinate KO (K00135) was not significantly different between seasons, but the dominant functional taxonomy shifted from Alphaproteobacteria in the summer stratified season to *Poseidoniia* (Thalassarchaeaceae) in the winter mixed season. In contrast, CE of the other genes was significantly enhanced or suppressed between summer stratified and winter mixed seasons (Fig 4B), meaning that the community’s total potential to produce or consume the exometabolite changed seasonally. Both observations could alter microbial production or consumption rates to produce apparent seasonal patterns of DOM features. Changes in rates could result from differences in enzymatic efficiency associated with the taxonomic shifts, a change in the total number of taxa capable of interacting with the exometabolite, or shifts in environmental conditions that induce changes in the regulation of reactions. These results emphasize that observations of labile molecules in the environment are the result of compounding mechanisms with different timescales, where turnover flux occurs on a timescale of days but also changes seasonally based on community structure.

Between years, we observed significant taxonomic variability of ASVs across our time-series (Fig 3), but we also identified core genes present in all historical metagenomes. The roles of microbial redundancy and diversity have been widely investigated (16, 41, 77). This study provides an additional lens to understand the resulting impacts on the exometabolome. At BATS, trehalose-metabolizing genes exhibited a specific-type redundancy, meaning trehalose utilization was limited to a narrow portion of the total community (83). In comparison, succinate metabolism was observed more globally across the microbial community and thus displayed a broad-type redundancy, which reflects its role in common microbial metabolisms. While these differences in utilization were expected based on previous literature, ecological theory would suggest that the lower functional redundancy of trehalose enzymes makes this metabolite more susceptible to variability when microbial community changes occur (77, 79). And yet, despite differences in functional redundancies, both succinate and trehalose exhibited similar seasonal patterns in the surface ocean that remained consistent across all years of the time-series (Fig 2). We suggest that as long as an undefined minimum threshold of functional redundancy is met, the composition of the seasonal exometabolome will remain consistent. Based on the time-series and metagenomic analyses, it appears that the role of a metabolite in reactions encoded by core genes of a microbial community is more important than the degree of functional redundancy. Even if a core gene is utilized only by a subset of community members, the associated metabolite will still be regularly exchanged through the labile DOM-microbe network.

## Conclusions

Understanding how the reservoir of marine microbial diversity translates into a similarly diverse pool of DOM molecules is a critical knowledge gap in our understanding of carbon cycling. In an environment that has already experienced 1.2°C of warming (84), resolving these baseline processes is essential in order to predict future changes in the ocean’s organic carbon cycle. We demonstrate that the metabolic functions, rather than taxonomic identity, of microbial communities are greater predictors of exometabolome composition.

The untargeted exometabolome time-series at BATS provided some of the first insights into the variability of DOM molecules on seasonal and interannual timescales, in parallel with the microbial community. Despite similar complexities with respect to composition, we found that the mechanisms responsible for driving bacterioplankton taxonomy and DOM molecules should be expected to differ. The results of our analyses are consistent with the perspective that many metabolic functions are shared across diverse, phylogenetically related taxa and ecological concepts such as Hubbell’s neutral theory, which predicts variation in species with no corresponding variation in metabolic function (85). This work highlights that in order to predict labile DOM flux, future models should focus on incorporating core metabolic pathways that are required for community function by either all or a portion of the microbial community.

Significant taxonomic variability was detected in the bacterioplankton community at BATS during the three-year time-series, but the changes were not enough to influence the composition of the resulting DOM biogeochemistry. This buffer of functional redundancy overlaid on taxonomic variability will play an important role in future oceans. How much can microbial taxonomy change, though, before the presence of these core metabolisms is altered? As anthropogenic carbon emissions alter the ocean’s temperature, pH, and nutrient supplies, microbial communities will shift and evolve in response, and, in some cases, may do so abruptly, which will inevitably have implications for DOM biogeochemistry (86–90). This work presents a major advance in our understanding of variability and composition of the individual molecules comprising DOM, as well as important avenues of research for predicting the resulting carbon flux. The seasonal patterns of DOM features represent snapshots of standing stocks, and future studies that emphasize rate measurements will be essential. Continuing to resolve the influences of the microbial loop’s functional redundancy and core metabolisms on DOM biogeochemistry is critical for predicting changes to the ecosystem function of heterotrophic carbon remineralization in future oceans.

## Materials and Methods

### Exometabolome sample collection and extraction

Samples were collected aboard the R/V *Atlantic Explorer* bi-monthly from fixed depths (1 m, 40 m, 120 m, 200 m) at or in the vicinity of the Bermuda Atlantic Time-series Study (BATS) site from July 2016 to July 2019. During July field campaigns, samples were collected from every sampling depth every 6 hours for 72 hours. During all other sampling events, one sample per depth was collected primarily between the hours of 05:00 and 10:00 local time. Samples were contextualized by a physical framework that defines the state of the water column across the seasonal cycle at BATS (Fig 1A) (31, 58). All major seasons were sampled every year, with the exception of the spring transition, which is a short-lived period and was missed in 2017 when it likely occurred between our sampling in April and May. 4L of whole seawater was filtered through a 47mm 0.2µm Omnipore PTFE filter (Millipore, Burlington MA, USA) using a peristaltic pump as described previously (91). 4L of onboard Milli-Q water was filtered in the same manner for process blanks. The filtrate was acidified to a pH of 2-3 with OmniTrace HCl (ThermoFisher Scientific, Waltham, MA, USA) and extracted via solid phase extraction with styrene-divinylbenzene polymer columns (1g Bond Elut PPL, Agilent, Santa Clara, CA, USA) as described previously (54, 92). Sample elutions were evaporated to near dryness and reconstituted in Milli-Q water with 22 stable isotope labeled internal injection standards (Table S4). A pooled sample was created with an aliquot of every sample.

### UHPLC-ESI-MSMS, exometabolite feature processing, and data filtering

The sample set (n = 374) was randomized across five batches. The pooled sample was used for column conditioning and was also injected after every 5 samples and at the end of each sequence, followed by process blanks and Milli-Q blanks. Batches were run in both positive and negative ionization mode. Chromatography was performed as previously described (60, 91) using an ultrahigh-performance liquid chromatography system (Vanquish UHPLC, Thermo Scientific) coupled with an Orbitrap Fusion Lumos Tribrid mass spectrometer (Thermo Fisher Scientific). Detailed instrument parameters are provided in the Supplement.

Raw data files were converted to mzML format using msConvert (93) and transferred to a high-performance computing cluster for processing with R (v 4.0.1). XCMS (v 3.10.2) was used for peak picking each sample and grouping shared peaks into a single feature (94). XCMS parameters and workflow are described in the Supplement. MS1 features were defined by a unique mass-to-charge ratio and retention time. The XCMS analysis yielded a table of MS1 feature intensities in each sample. Presented intensities are unitless as this integration reflects an integration of all ion counts associated with a given feature’s mass-to-charge ratio bounded by the retention time window. CAMERA was used to identify and filter isotopologues and adducts (95). Features were further filtered for peak quality, blank contaminants, inter-batch variability, and detection in the samples as described in the Supplement. Feature intensities were batch corrected using the BatchCorrMetabolomics package (v 0.1.14) with a robust least-squares regression (96). Well-behaved injection standards exhibited an RSD <20% across all injections after batch correction (Table S4), which is an acceptable threshold for large-scale metabolomics experiments (97). XCMS was also used to produce .mgf files (consensus spectra and maximum total ion current spectra) and abundance tables, which were submitted to the Global Natural Products Social Molecular Networking infrastructure for feature-based molecular networking (98). The GNPS results putatively identified the four exometabolites presented herein, which were then further validated with authentic standards (Table S1).

### Microbial community, 16S rRNA amplicon sequencing, and data filtering

Samples for 16S V1-V2 amplicon sequence variants (ASVs) were collected as described in Liu et al. (2022). Only samples collected at 1 m, 40 m, 120 m, and 200 m were presented. Briefly, 4L of seawater were filtered onto 0.2 μm Sterivex and stored at −80**°**C. DNA was extracted with a phenol-chloroform protocol (67). V1-V2 16S rRNA hypervariable region was amplified with primers 27F (5′-AGAGTTTGATCNTGGCTCAG-3′) and 338RPL (5′-GCWGCCWCCCGTAGGWGT-3′). Amplicon libraries were built using the Nextera XT Index Kit (Illumina Inc.) and sequenced using the Illumina MiSeq platform (reagent kit v.2; 2×250 PE) at the Center for Quantitative Life Sciences (CQLS), Oregon State University. Raw amplicon datasets were processed as in Bolaños et al. (2022) using Dada2 v1.18 (99) with the following filtering parameters: maxEE=(2,2), truncQ=2, minLen=190, truncLen= 220,190, maxN=0. Samples from the same sequencing run were processed together to accurately estimate the error frequency. Potential chimeras were removed with the removeChimeraDenovo command. Taxonomic assignment was performed with the assignTaxonomy command and the Silva non redundant database V.123 (100). Generated ASV and taxonomic tables were analyzed using phyloseq v1.34 (101). ASVs were presented as relative abundances, normalized to the total counts of all ASVs in a respective sample. ASVs were required to be detected in ≥ 5% of all samples. This yielded an ASV table with n = 3158 taxa. The terminal node collapse of the ASVs was conducted via PhyloAssigner with a global reference tree, resulting in a table of n = 1806 taxa (82).

### Wavelet analysis

Wavelet analysis was used to decompose the exometabolome and ASV time-series using the R package WaveletComp (34) (Fig S1). Wavelet analysis requires a uniform grid. Most of the time-series was sampled in odd months, except for samples collected in April. We interpolated between months to create a monthly time-series that allowed us to utilize the April data. This also avoided any distortion to the wavelet analysis which is sensitive to time-series length. In months where more than one sample was collected, we used the average feature intensity as the representative value. Significance was assessed with the null hypothesis of white noise and 1000 permutations were calculated for each time-series. Similar trends were observed for the exometabolome in both ionization modes, and therefore only positive mode results were discussed.

### Metabolic redundancy and functional taxonomy

HMMER (v 3.3.1, hmmer.org) searches were conducted with HMM profiles previously created by KofamScan (102). KO numbers were collected based on analysis of KEGG Pathways (103) to find key enzymatic reactions required to conduct pathways that result in the production or consumption of trehalose and succinate (Table S*2*). Multiple KOs can encode for the same metabolic transformation, and for brevity we present the most redundant KO only (Fig 4). In addition, single copy marker genes (SCMG) were also searched to estimate sample richness (K01409, K01869, K01873, K01875, K01883, K01887, K01889, K03106, K03110, K06942). Publicly available surface sample metagenomes collected at BATS were queried from 1997 – 2019, though with non-uniform sampling (Table S3). Metabolic KO HMM results were filtered with an e-value of 1 x 10^-10^, and SCMG KOs were filtered based on threshold scores defined by KofamScan. Samples richness was calculated based on the number of contigs encoding a SCMG. The taxonomy of metabolic KO genes was assigned using the contig level taxonomy annotations from MDMcleaner (v 0.8.2) ‘clean’ output with ‘—fast_run’ settings (104). Presence was calculated as the number of contigs assigned to a metabolic KO. Traditional metrics of functional redundancy, which are based on niche space, are not easily translated for microbial communities. Here we calculated the metric of contribution evenness as an estimate of metabolic redundancy (79).

### Data presentation

All figures were created with ggplot2 (v 3.4.3) and curated with Inkscape (v 1.2.2).

## Data availability statement

Metabolomics data, including raw files, mzML files, and feature tables, are deposited at MetaboLights under study accession number MTBLS5228. 16S amplicon sequences are deposited in the National Center for Biotechnology Information (NCBI) Sequence Read Archive (SRA) under project number PRJNA769790. Publicly available metagenomes were accessed from NCBI SRA project number PRJNA385855 (105) and newly deposited historical metagenomes from NCBI SRA project number PRJNA769790. CTD data are deposited in the Biological and Chemical Oceanography Data Management Office (BCO-DMO) at http://lod.bco-dmo.org/id/dataset/861266 for BIOS-SCOPE cruises, and at http://lod.bco-dmo.org/id/dataset/3782 for BATS cruises.

Code for processing the raw exometabolome data is available at https://github.com/KujawinskiLaboratory/UntargCode. Code for processing the raw amplicon data is available at https://github.com/lbolanos32/NAAMES_2020. Code for PhyloAssigner, analyzing the time-series, querying the metagenomes, and calculating metabolic redundancy is available under git project https://github.com/BIOS-SCOPE/FunctionalRedundancy

## Acknowledgements

We thank Rod Johnson, the BATS technical team, especially Julia Matheson and Paul Lethaby, the BIOS-SCOPE team for their efforts during sample collection, the MAGIC lab for hydrographic data, and the R/V *Atlantic Explorer* officers, technicians, and crew members. We greatly appreciate work by many others to develop and share the publicly available code and data utilized in this study. We thank Georg Pohnert and Muhaiminatul Azizah for sharing the gonyol reference standard. We also thank anonymous peer reviewers for helpful feedback, as well as Brianna Garcia, Anya Brown, and Arianna Krinos. This project was funded by the Simons Foundation International’s BIOS-SCOPE program. McParland was funded by the Woods Hole Oceanographic Institution Postdoctoral Scholar program and the Simons Postdoctoral Fellowship in Marine Microbial Ecology. This is the NSF Center for Chemical Currencies of a Microbial Planet (C- CoMP) publication #037.

## Supplement

### Supplemental Text 1: Methods

#### UHPLC-ESI-MS/MS performance

Separation was performed with a reverse phase Waters Acquity HSS T3 column (2.1 x 100 mm, 1.8 μm), equipped with a Vanguard pre-column. Column temperature was held at 40°C. The column was eluted at 0.5 ml/min with a combination of solvents: A) 0.1% formic acid in water and B) 0.1% formic acid in acetonitrile. The chromatographic gradient was as follows: 1% B (1 min), 15% B (1-3 min), 50% B (3-6 min), 95% B (6-9 min), 95% B (10 min). The column was washed and re-equilibrated with 1% B (2 min) between injections. The autosampler was set to 4°C and injection volumes were 5 µl. The electrospray voltage was set to 2600 V for negative mode and 3600 V for positive mode. The setting for source sheath gas was 55 and auxiliary gas was 20 (arbitrary units). The heated capillary temperature was 350°C and the vaporizer temperature was 400°C. MS data were collected in the Orbitrap analyzer with a mass resolution of 120,000 FWHM at m/z 200. The automatic gain control (AGC) target was 4e^5^, the maximum injection time was 50 ms, and the scan range was 100 – 1000 m/z. Internal mass calibration of the Orbitrap analyzer was used to improve mass accuracy of the MS scan. Data-dependent MS/MS data were acquired in the Orbitrap analyzer using higher energy collisional dissociation (HCD) with a normalized collision energy of 35% and with mass resolution of 7500. The AGC target value for fragmentation spectra was 5e^4^ and the intensity threshold was 2e^4^. Cycle time was set at 0.6 s. Precursor selection was performed within the quadrupole with a 1 m/z isolation window. Dynamic exclusion was enabled, with 3s exclusion duration after n=1. All data were collected in profile mode. Raw data files were converted to mzML format using msConvert (1).

Large LC-MS/MS experiments are prone to retention time drift, contamination, and carry over between samples (2). To mitigate these factors LC-MS sequences were limited to 105 injections (∼18 hrs), the internal mass calibration was enabled (equivalent to a lock mass correction), the column was re-equilibrated at the beginning of each batch, sample order was randomized, the ESI probe was cleaned between batches, multiple stable-isotope labeled internal injection standards were added to all samples, and a pool QC sample was run after every n = 5 samples. All of these efforts were successful in mitigating unwanted variation (see Supplemental Text 2), with the exception of our QC sample. After running all batches, it was discovered that the pool sample was sub-sampled too many times and thus created a linear decrease in the TIC of these injections over time across each batch that could not be compared. However, one pool sample per batch was aliquoted, and thus variability could be calculated within the QC by comparing the first injection of each batch (n = 5).

#### XCMS and CAMERA workflow

Peak-picking was performed using the CentWave algorithm with the following parameters: noise = 100, peak-width = 3-14, ppm = 15, prescan = 3, preintensity = 5e4, snthresh = 0, integrate = 2, mzdiff = −0.005, extendLengthMSW = TRUE, fitgauss = FALSE, firstBaselineCheck = FALSE. Replicate picked peaks were merged with refineChromPeaks (MergeNeighboringPeaks Param: expandRt = 0, expandMz = 0, ppm = 5, minProp = 0.75). Peaks were filtered based on peak quality by requiring a peakwidth less than 15 seconds and with a custom R script based on Gaussian fits (correlation value > 0.6 and a p-value < 0.075). Retention times were adjusted using Orbiwarp (binSize = 0.1) based on the center sample (3). Correspondence (bw = 0.7, binSize = 0.0005) between the peaks was conducted using the peak density method (4). As every effort was made to optimize accurate parameters for peak picking, alignment, and correspondence based on internal injection standards and manually checked DOM features, we did not utilize the fillChromPeaks here as we found it primarily resulted in the integration of noise. Feature values were integrated by the ‘maxint’ method. CAMERA was performed to identify isotopes and adducts by grouping features based on retention time to create pseudospectra (perfwhm = 0.5), identifying ^13^C isotopologues (ppm = 3, mzabs = 0.01), and grouping based on correlations of intensity, extracted ion chromatograms, and isotopes (corr_eic_th =0.9, cor_exp_th=0.8, pval=0.05) (5).

#### Feature filtering

The XCMS and CAMERA workflow resulted in n = 153,360 features in positive mode and n = 117,079 features in negative mode. By optimizing XCMS performance to maximize peak picking, a majority of these resulting MS1 features were noise and we therefore performed stringent best-practices for feature filtration. Features were filtered based on results from CAMERA to remove identified isotopologues and adducts (5). Features were filtered using Milli-Q and process blanks using a data-adaptive method (6). The mean log abundance across samples and blanks was calculated for each feature, and subsequently binned into 20, 40, 60, and 80^th^ quantiles. For each bin, a threshold was calculated based on the 25^th^ quartile of the difference between the mean log abundances of samples and blanks that were less than 0. The difference for all features in a given bin were required to be greater than the absolute value of this threshold. Features were filtered to require their grouped peaks to have a range in median retention times of less than 5 seconds. Features were filtered to require their detection in >50% of all samples. If a feature was detected in the pool sample, it was required to have a relative standard deviation < 30% as calculated based on the intensity across the first pool sample injected in each batch (n = 5). The filtered features total 4% of the original features output by our XCMS workflow. The remaining features represent, to the best of our ability, unique molecules (defined by a m/z and retention time), but these datasets will always contain undistinguishable isomers, adducts, and isotopologues (7, 8).

#### Metabolite identification

All four putative identifications had m/z matches to reference masses within ± 1ppm (Fig S6 - Fig S9). The identifications were originally made by GNPS and subsequently confirmed with authentic standards when possible. Based on confidence levels defined by the Metabolomics Standards Initiative (9), succinate, trehalose, and gonyol were identified to the highest level possible (Level 1) using standards analyzed by the same analytical platform used to analyze the untargeted exometabolome. Glucose 6-sulfate was identified to the second highest confidence level (Level 2) as, to the best of our knowledge, an authentic standard for this compound does not exist. The putative identification was made based on a match to a reference spectrum of the almost identical compound, glucose 6-phosphate. However, the exact mass difference between the two different precursor masses (0.009 m/z) is equal to the expected mass difference between glucose 6-sulfate and glucose 6-phosphate (0.0095 m/z). Additionally, the MS2 spectrum supports this identification. A dominant MS2 fragment was m/z 96.959 (HSO_4-_), in comparison a phosphate containing fragment would have a mass of m/z 96.969 (H_2_PO_4_). Although the exact masses support the presence of the sulfate group, we cannot rule out that the putatively identified glucose 6-sulfate could instead be the isomer galactose 6-sulfate.

### Supplemental Text 2: Unwanted variability in untargeted exometabolomics

Here we discuss potential sources of unwanted variability due to instrumentation and computational processing and estimate their presence in this dataset. To minimize variability induced by changes in instrument performance between batches, we applied a robust least-squares regression to each DOM feature (10). This regression shifts the mean of each batch so that intra-batch variability is maintained but is centered on a common mean. QC pool samples in large-scale metabolomics experiments are expected to exhibit a variation of <20% (relative standard deviation) (2). This value will encompass any inter-batch variability not removed by the batch correction, intra-batch variability from instrument performance, and variability induced from computational preprocessing (10–12). To estimate the amount of unwanted variation in our sample set we utilized both QC pool samples and stable-isotope labeled injection standards (added after solid phase extraction). As described above, we could not use all of our QC pool samples to quantify unwanted variability. However, based on the first injection of the QC pool sample for each batch we filtered features by requiring <20% inter-batch variability. A total of n = 22 stable isotope labeled internal injection standards were added to all samples, of which n = 12 were expected to ionize in positive mode. In order to be used for calculating unwanted variability, “well-behaved” injection standards were required to: 1. exhibit good quality peak shapes based on visual inspections, 2. exhibit peak heights above limits of detection, 3. display stable retention times, and 4. be detectable by XCMS. A total of n = 8 injection standards behaved well in positive ionization mode. All of these injection standards had a relative standard deviation ≤ 20%. It is noteworthy that this variation is calculated across all samples (1-200m, and all seasons), meaning that any changes in ionization due to changes in bulk DOM across the time-series did not significantly alter the behavior of the injection standards above acceptable thresholds. Based on our analysis, we strongly support the inclusion of multiple stable-isotope labeled internal injection standards (13). The internal injection standards were used to quantify unwanted variability, but were also essential for optimizing XCMS performance.

### Supplemental Text 3: Putatively identified exometabolites’ role in the ocean

Here we further discuss the potential role of putatively identified exometabolites in the marine microbial loop. Gonyol is a reduced organic sulfur molecule similar in structure to the well-known metabolite dimethylsulfoniopropionate (DMSP) (14). Gonyol is produced with taxonomic-specificity and can be degraded by marine bacteria, but the genetic pathways responsible for recycling this exometabolite are not yet known (15–17). As would be expected for a labile molecule, the first quantification of gonyol in the dissolved phase was found at low nM concentrations in the Pacific Ocean (18), and our time-series suggests that these concentrations would likely change seasonally. Glucose 6-sulfate is similar to the core metabolite glucose 6- phosphate, where the phosphate group is substituted for an oxidized sulfate group. To our knowledge, this is the first detection of glucose 6-sulfate in the oligotrophic ocean, and its potential sources or sinks remain an open question. Glucose 6-sulfate (or its isomer galactose 6-sulfate) could be a degradation product of presumably abundant, but poorly characterized, sulfated polysaccharides that comprise algal cell walls (19). Little is known about these large biopolymers in DOM, and most knowledge is derived from studies of macroalgae, which would include *Sargassum* at BATS (20). Nevertheless, microalgae and bacteria can also produce sulfated polysaccharides (21). A diverse suite of known sulfatases could be utilized to remove the oxidized sulfate group and consume the remaining monosaccharide (21–23). Many questions remain to be answered about these two organic sulfur molecules, particularly considering past work in phosphorus-limited waters of the North Atlantic Ocean that has demonstrated the substitution of sulfur into core biomolecules (24).

Trehalose and succinate are both metabolites for which little is known about their presence in marine DOM, but their genetic pathways are well-characterized. Trehalose is a sugar that can be easily routed to glycolysis after breaking the disaccharide bond, but has also been shown to be synthesized or retained as an osmolyte (25–28). Succinate is a dicarboxylic acid produced as an intermediate metabolic product of the citric acid cycle and glyoxylate pathway making it a key part of core catabolism and anabolism pathways in marine microbes (29–31).

## Supplementary Figure Captions

**Fig S1:**
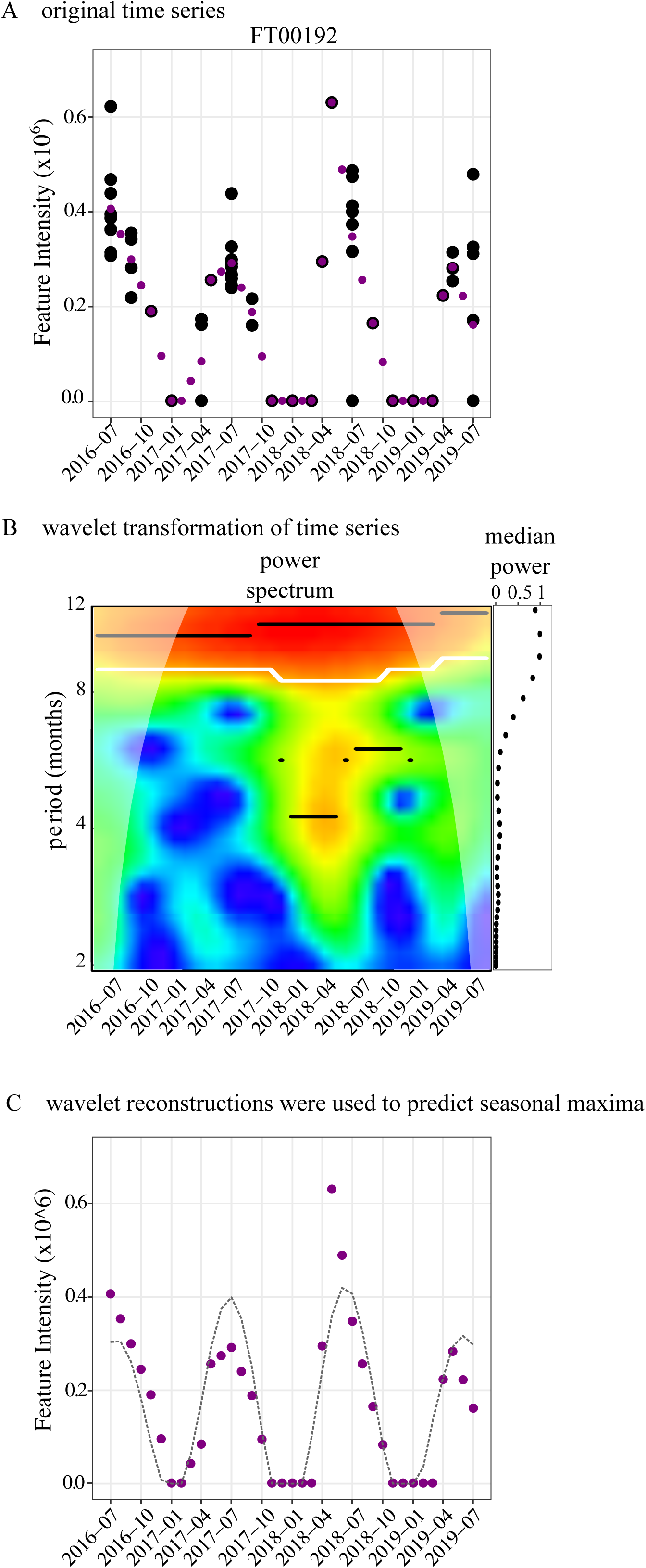
Example of the time-series wavelet analysis. (A) The original time-series (black) is almost uniformly sampled and contains multiple samples from the same month in some cases. The time-series was first transformed by averaging for months with multiple samples and interpolating between months to create a time-series with one sample per month (purple) (n = 37 samples; see Methods). (B) Wavelet analysis is used to detrend the time-series as reflected by the resulting power spectrum, where the calculated power (colorbar) for every sample (x-axis) is plotted as a function of every calculated period (1-12 months). The x-axis is the same as that in panel A. The side panel represents the median power for each period. A higher median power indicates a better wavelet fit. The highest median power was used to assign the dominant period of a time-series. In this example, the median power is highest for a period of ≥ 11 months and was therefore considered to be a seasonal time-series. (C) If the highest significant median power was ≥ 11 months, the wavelet was reconstructed (dashed black line) using a period of 12 months to predict the seasonal maximum. The month of the maximum value in the reconstructed time-series was used to assign the peak season.

**Fig S2:**
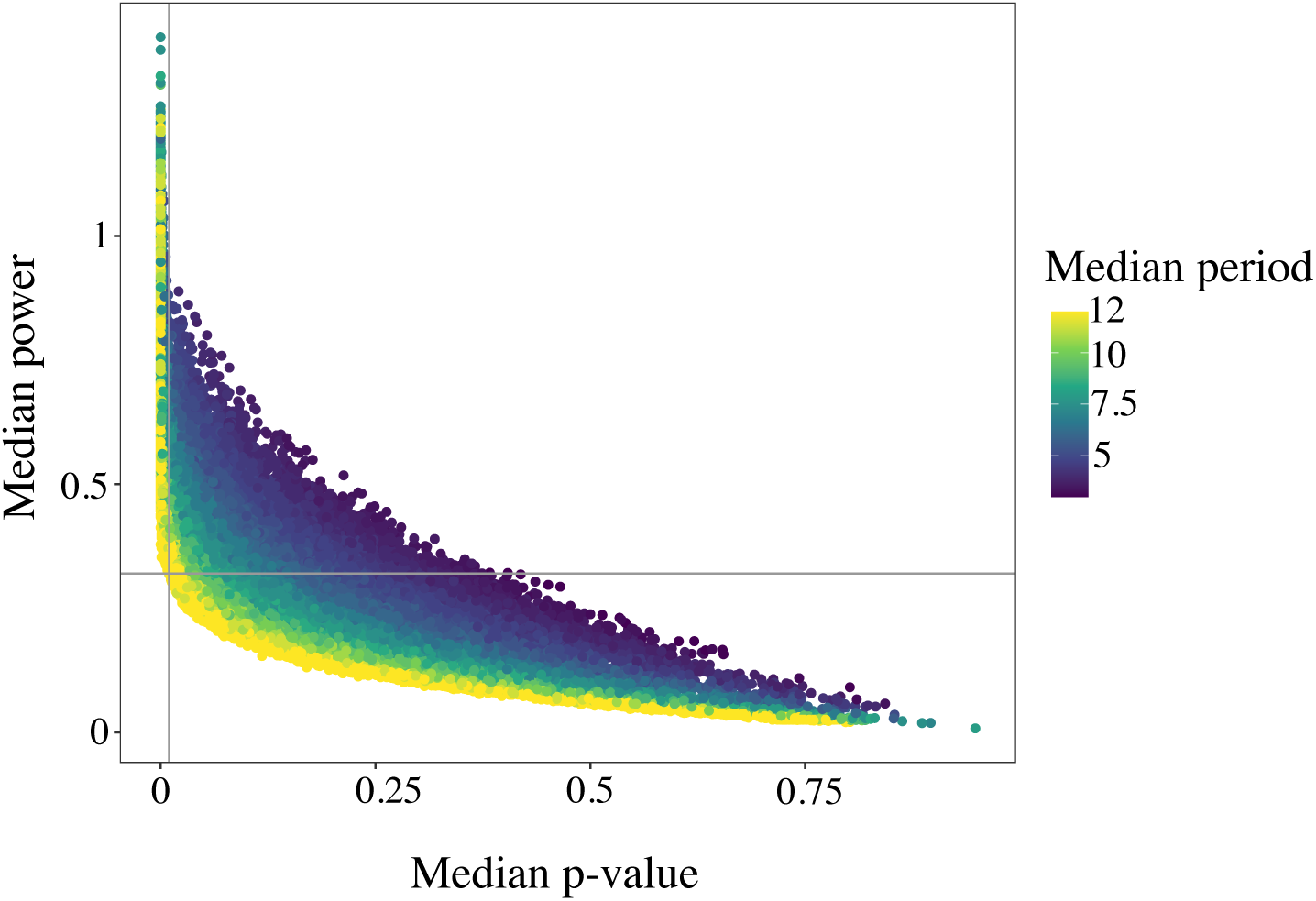
The median p-value and median power of all wavelets. Vertical grey line represents the median p-value cut-off of 0.01 and the horizontal grey lines represents the resulting minimum possible median power of 0.32. Color reflects the median period (1-12 months).

**Fig S3:**
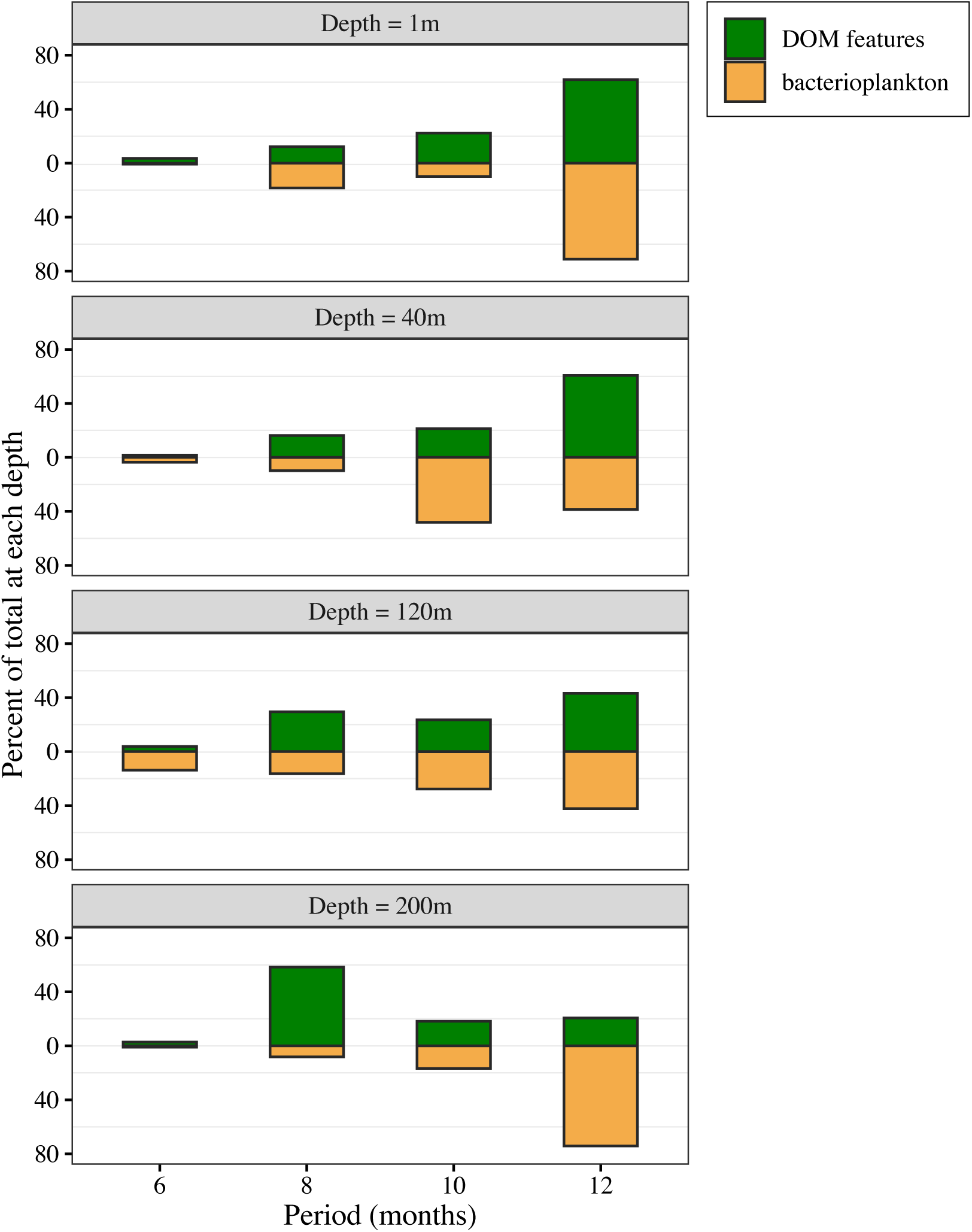
The percentage of all significant wavelets found in time-series with a dominant period of 5-12 months for DOM features (green) and bacterioplankton (yellow) at each sampling depth (1, 40, 120, and 200 m).

**Fig S4:**
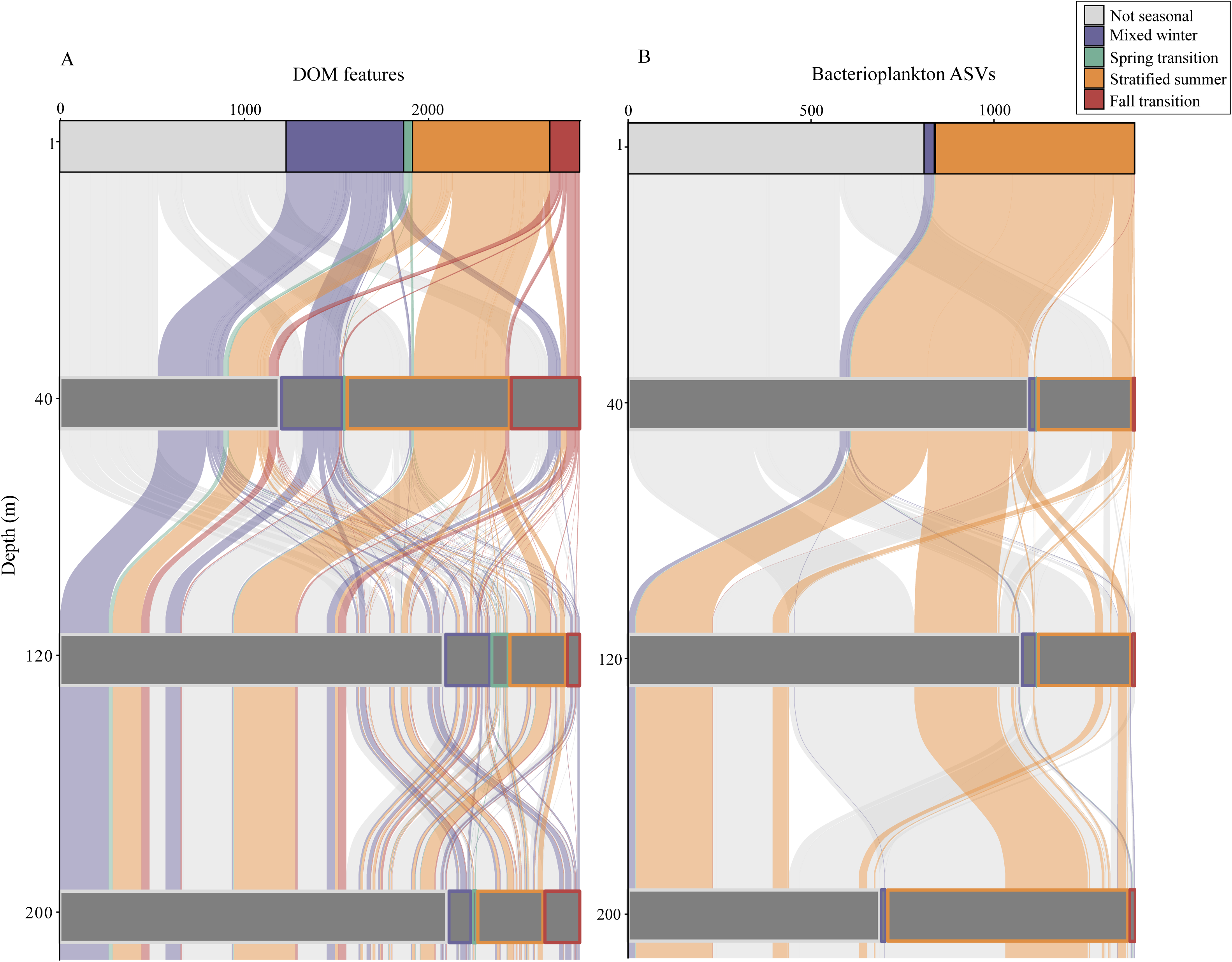
Alluvial plots depicting the connectivity of seasonal (A) DOM features and (B) bacterioplankton across sampling depths. Horizontal boxes represent the total number of seasonal DOM features or bacterioplankton, while box width and color reflect the number of features that peaked in a given season at the respective sampling depth. Grey represents a feature that is not seasonal at that depth but becomes seasonal at another depth. The ribbon colors track the connectivity of seasonal DOM features or bacterioplankton at 1 m through the water column.

**Fig S5:**
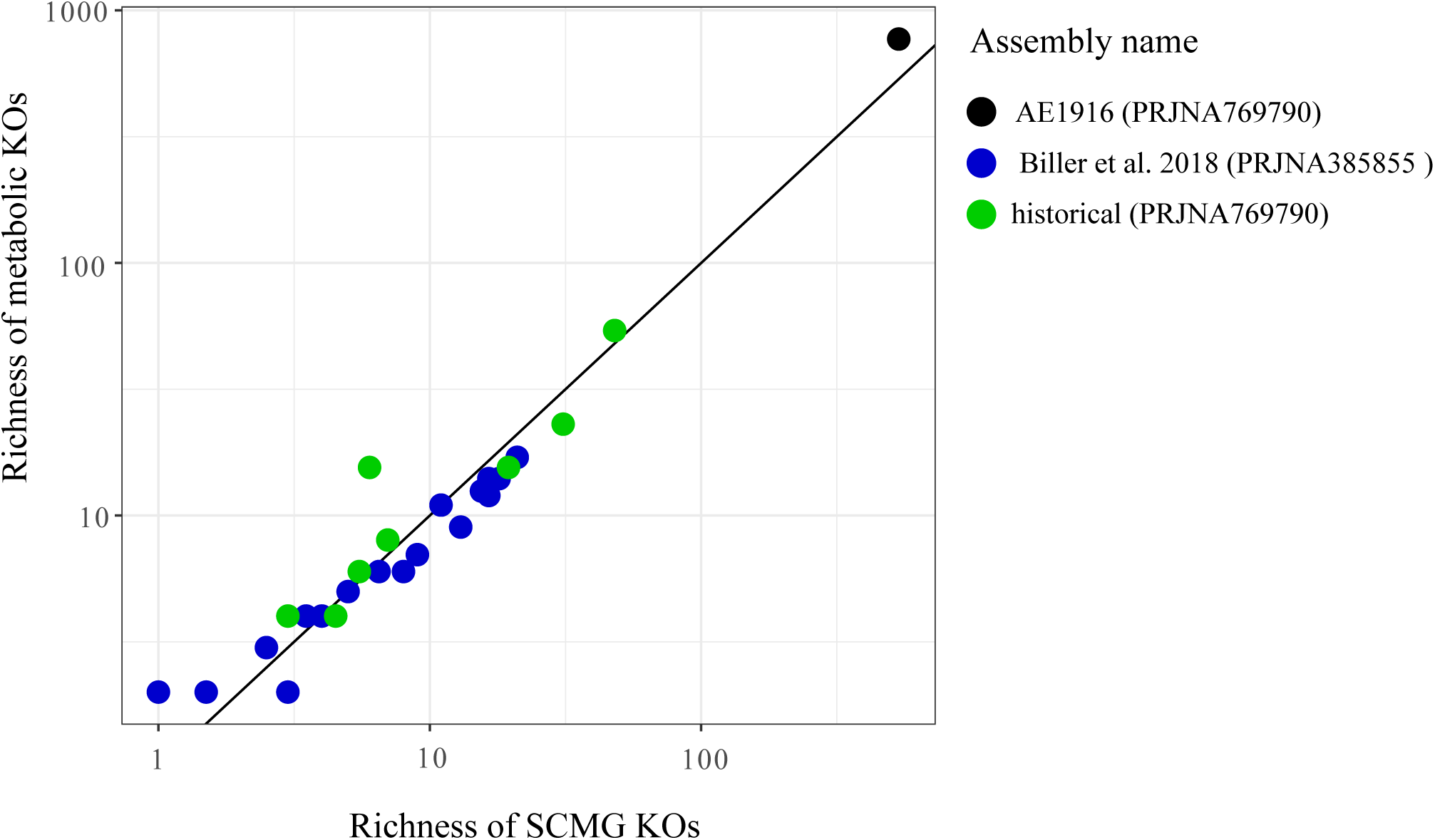
Median richness of all single copy marker gene (SCMG) KOs versus median richness of all metabolic functions (trehalose and succinate KOs) in all surface samples (n = 30) of each assembly queried. Color reflects the assembly name as defined in Table S3. The black line reflects a 1:1 relationship to demonstrate the linear relationship between the two groups of KOs queried, despite the order of magnitude differences in sequencing depth across the 20 years of metagenomic information.

**Fig S6:**
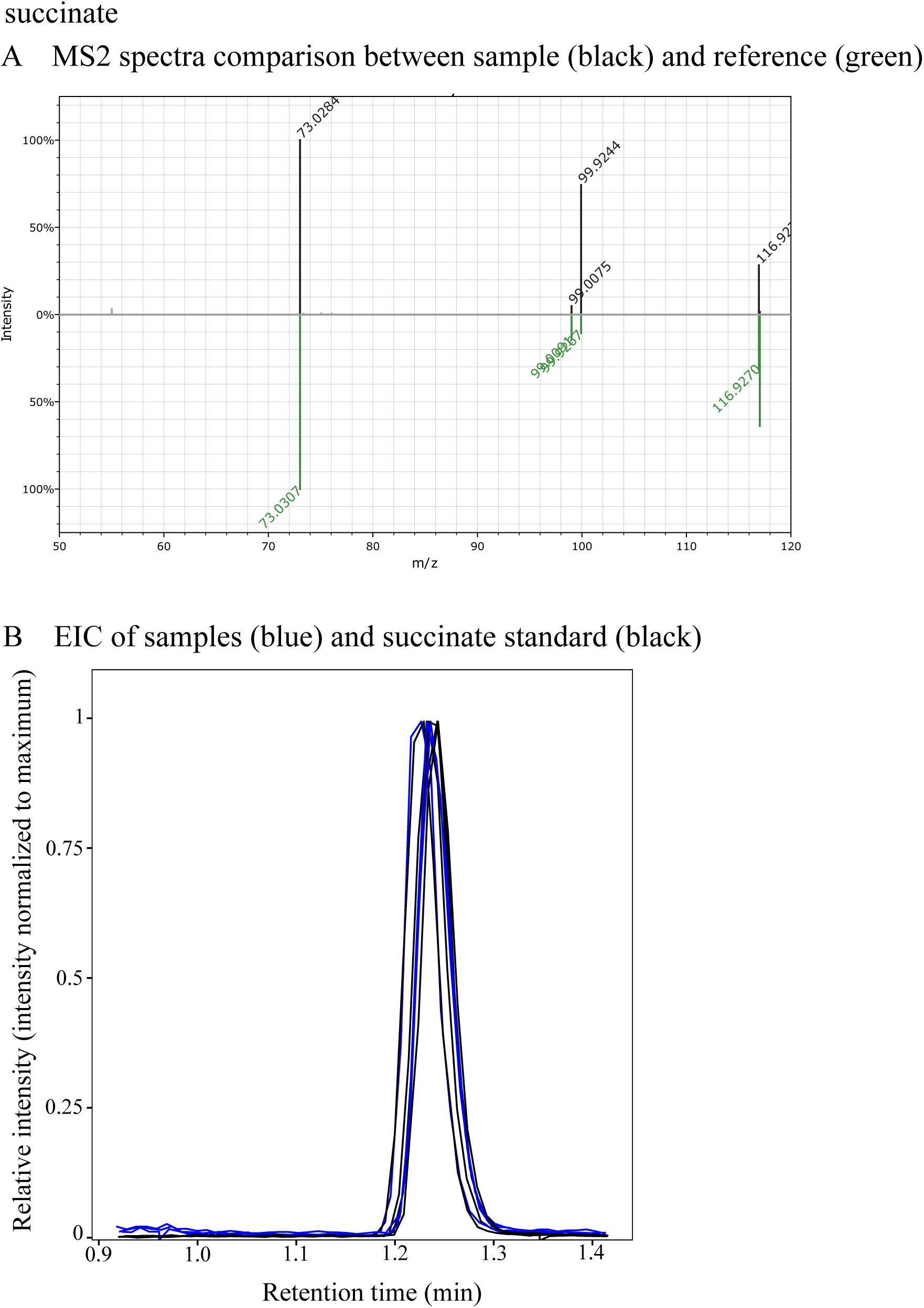
Succinate putative identification (Level 1). (A) Mirror plot of common MS2 fragments from samples (black) compared to the succinate reference spectrum in GNPS (green). (B) EIC of samples (blue) compared to succinate standard (black).

**Fig S7:**
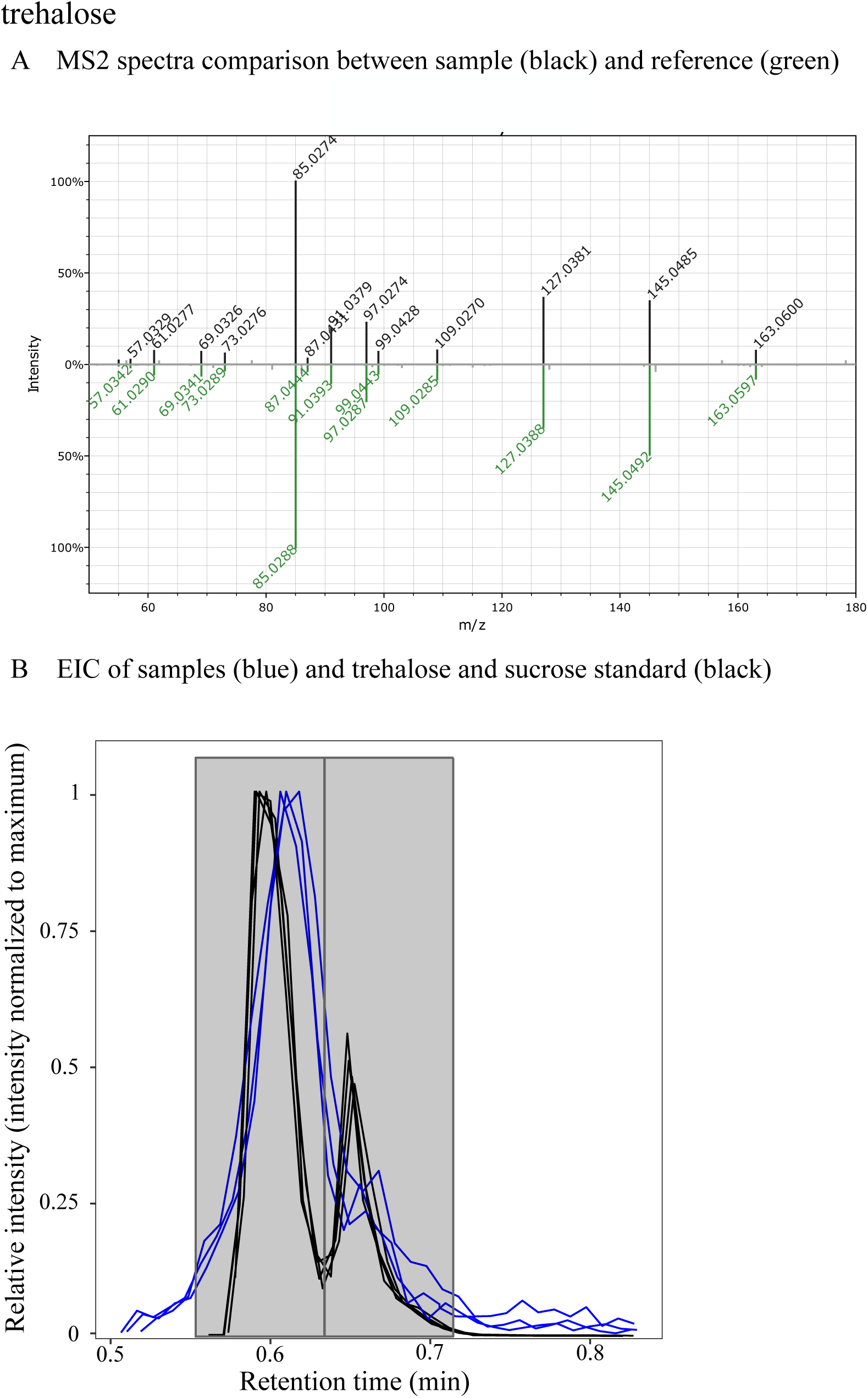
Trehalose putative identification (Level 1). (A) Mirror plot of common MS2 fragments from samples (black) compared to the trehalose reference spectrum in GNPS (green). (B) EIC of samples (blue) compared to trehalose and sucrose standards (black). The grey box highlights the chromatographic separation of trehalose and sucrose.

**Fig S8:**
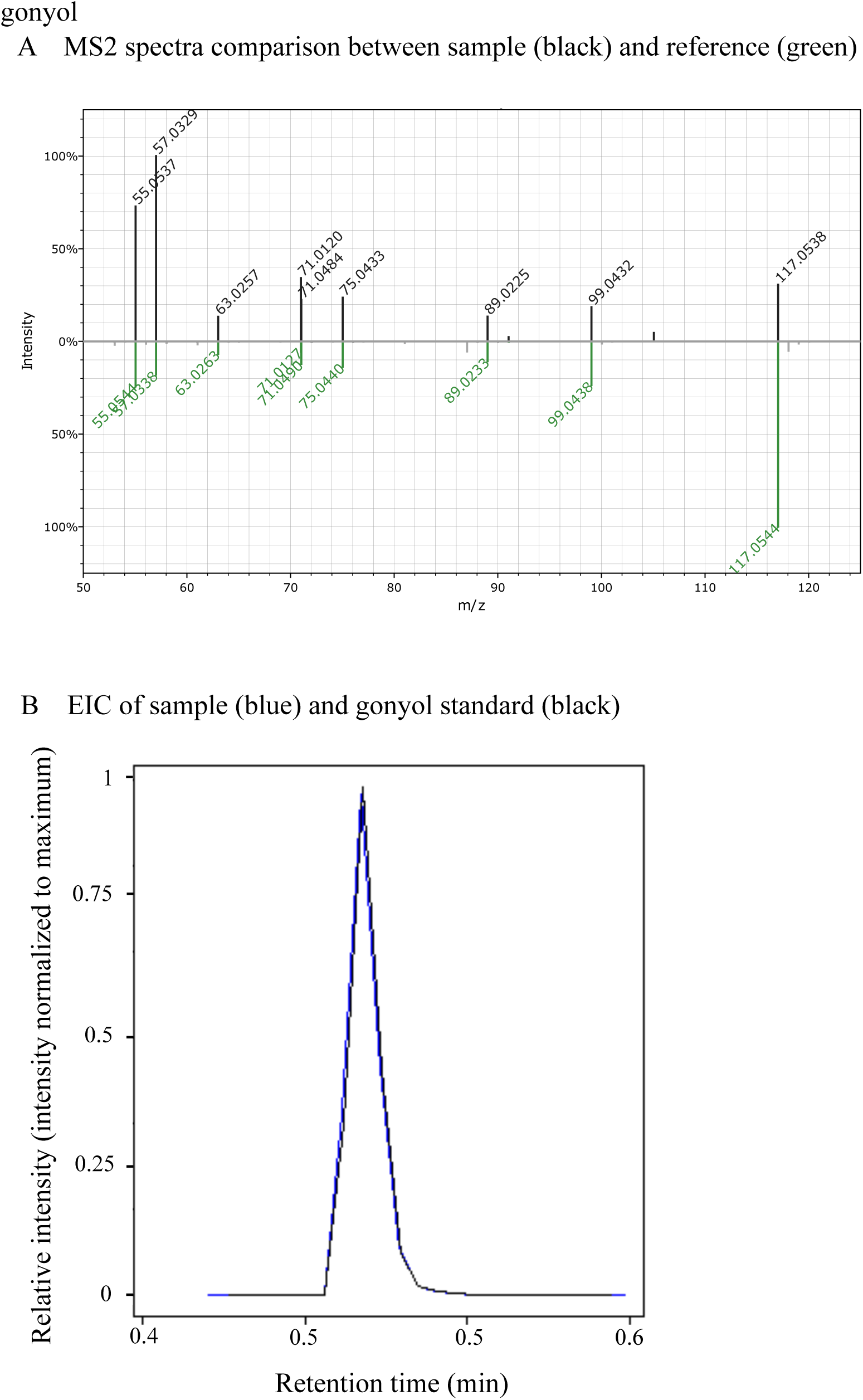
Gonyol putative identification (Level 1) (A). Mirror plot of common MS2 fragments from samples (black) compared to the gonyol reference spectrum in GNPS (green). (B) EIC of samples (blue compared to gonyol standard (black).

**Fig S9:**
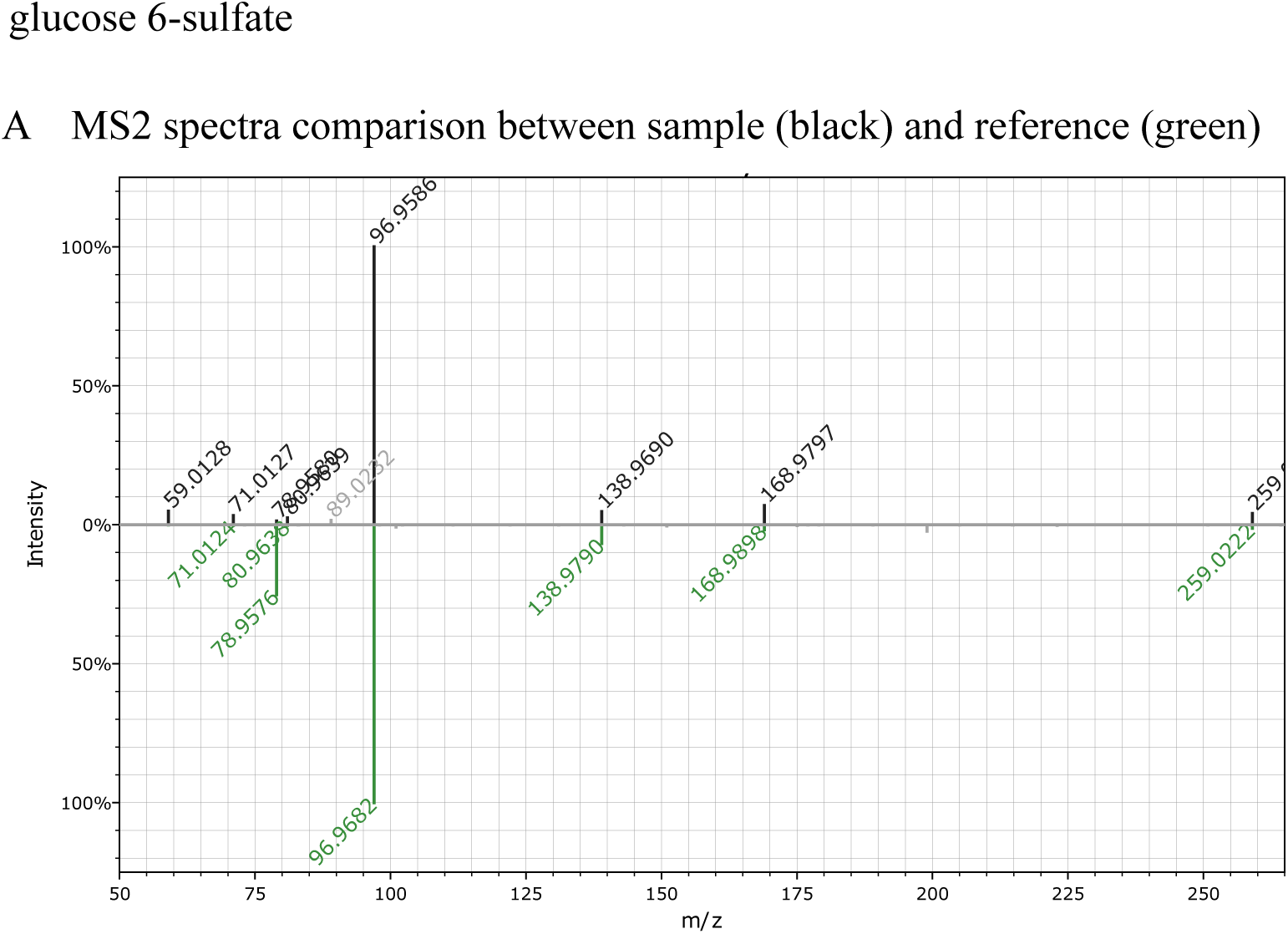
Glucose 6-sulfate (or galactose 6-sulfate) putative identification (Level 2). (A) Mirror plot of common MS2 fragments from samples (black) compared to the glucose 6-phosphate reference spectrum in GNPS (green).

## Supp Table Captions

**Table S1:**
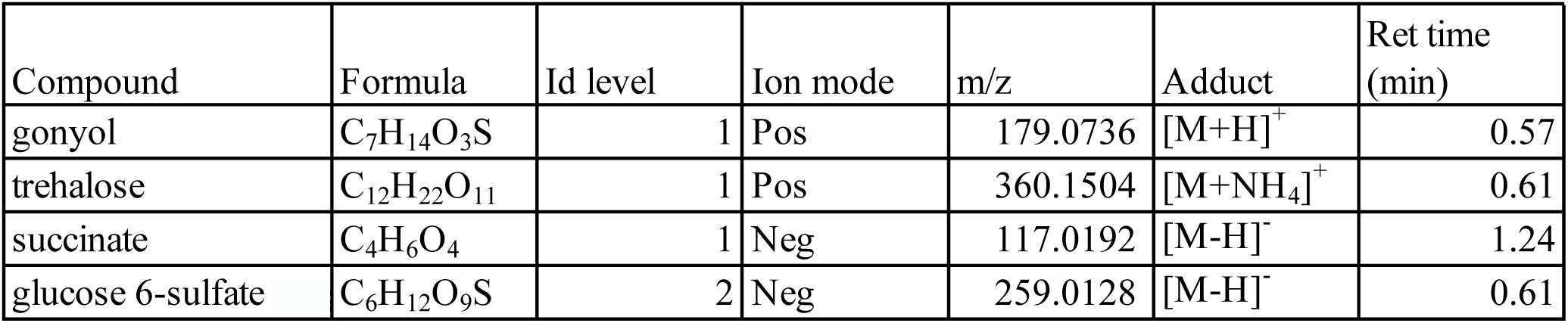
Molecular formula, identification level as defined by the Metabolomics Standards Initiative (9), ionization mode, detected m/z, adduct, retention time, and of putatively identified metabolites.

**Table S2:**
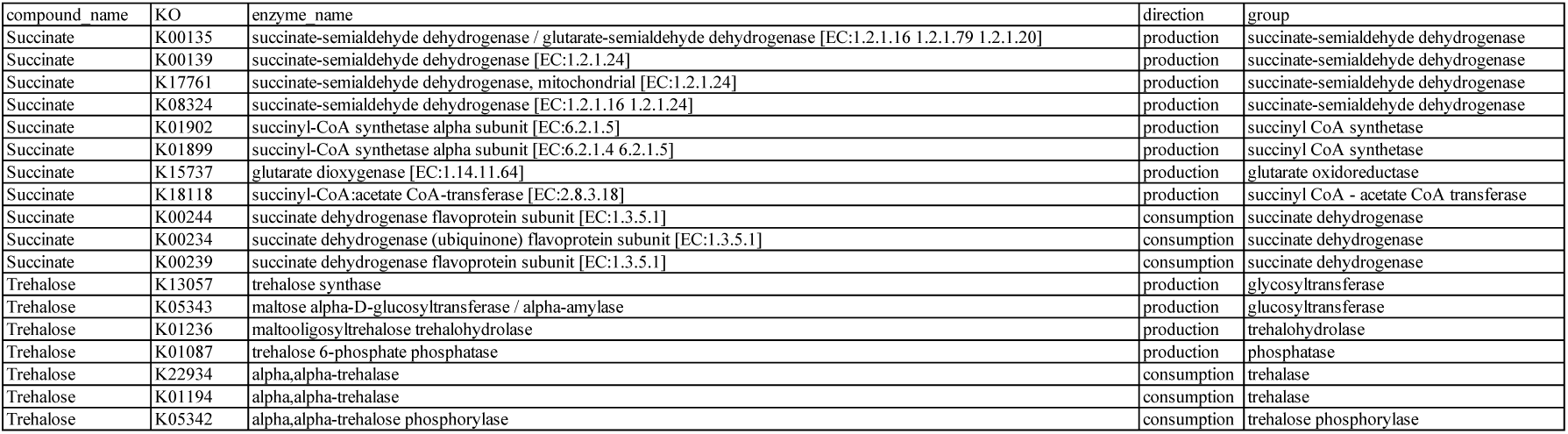
List of metabolic KOs queried in all surface metagenomes for trehalose and succinate production or consumption.

**Table S3:**
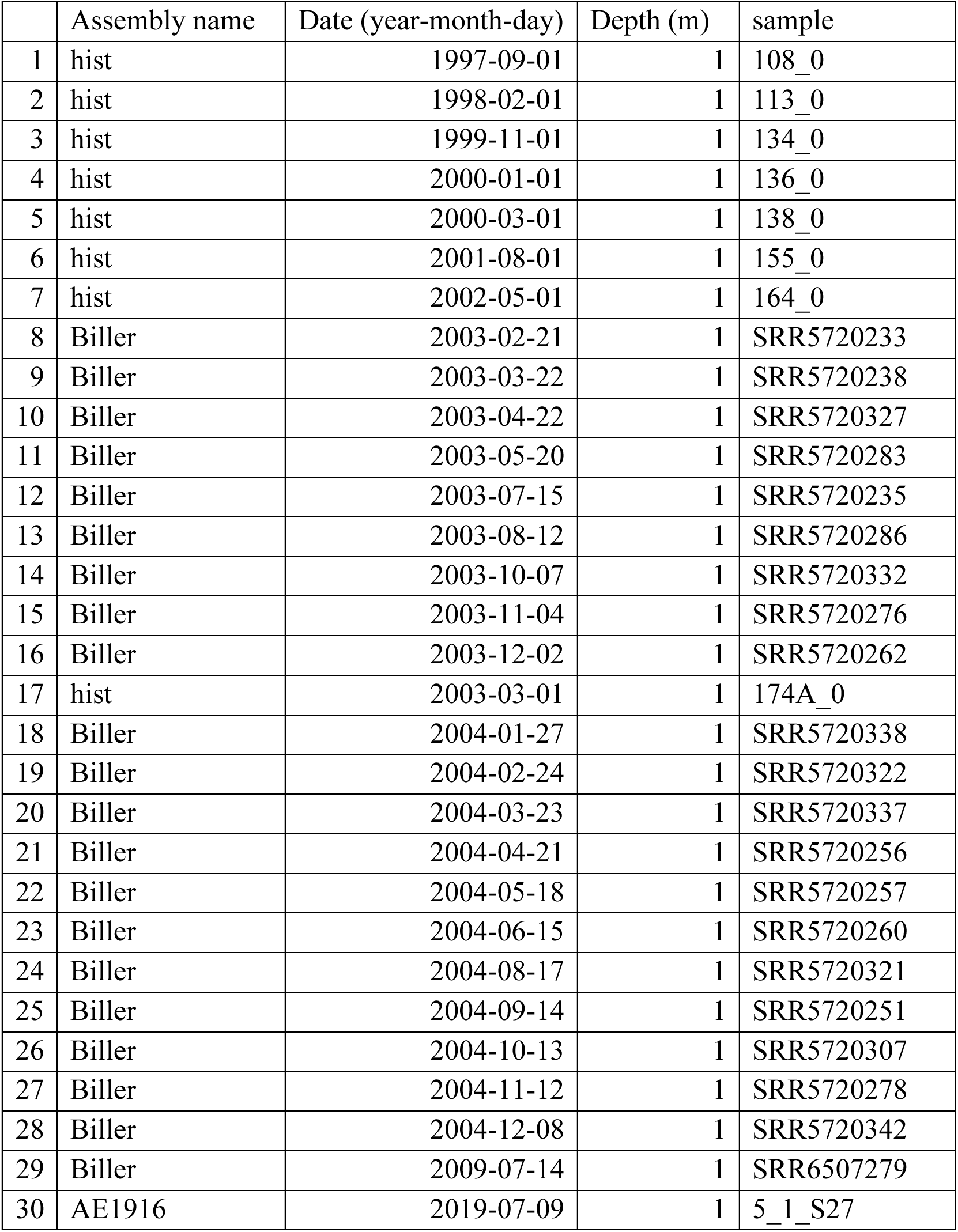
Metagenome assemblies queried for functional redundancy analyses.

**Table S4:**
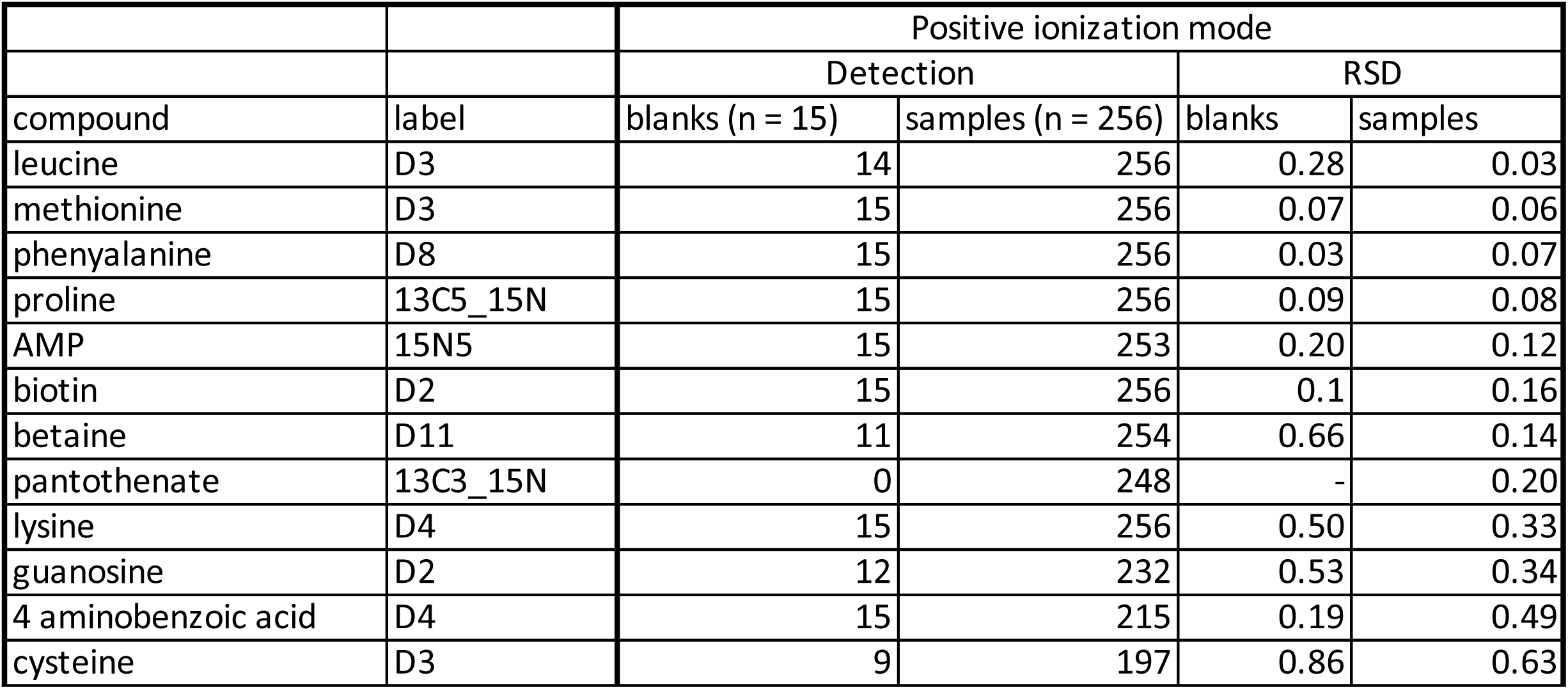
A mix of stable isotope labeled internal standards was added to every sample of the exometabolome. The label reflects which element was isotopically heavy. Ng/ml reflects the concentration added to each sample. Ion mode reflects which ionization mode the standard was detected in. The retention time reflects where the standard was detected in the chromatogram. The exact mass was calculated based on monoisotopic element composition. All standards were detected as either [M+H] or [M-H] adducts.

## References

1. C. A. Carlson, S. Liu, B. M. Stephens, C. J. English, “DOM production, removal, and transformation processes in marine systems” in Biogeochemistry of Marine Dissolved Organic Matter, Third, D. A. Hansell, C. A. Carlson, Eds. (Elsevier, 2024), pp. 137–246.

2. Y. Wang, et al., Linking Microbial Population Succession and DOM Molecular Changes in *Synechococcus*-Derived Organic Matter Addition Incubation. Microbiol Spectr 10 (2022).

3. E. K. Wear, C. A. Carlson, M. J. Church, Bacterioplankton metabolism of phytoplankton lysates across a cyclone-anticyclone eddy dipole impacts the cycling of semi-labile organic matter in the photic zone. Limnol Oceanogr 65, 1608–1622 (2020).

4. C. A. Carlson, et al., Interactions among dissolved organic carbon, microbial processes, and community structure in the mesopelagic zone of the northwestern Sargasso Sea. Limnol Oceanogr 49, 1073–1083 (2004).

5. F. Vincent, et al., Viral infection switches the balance between bacterial and eukaryotic recyclers of organic matter during coccolithophore blooms. Nat Commun 14, 510–17 (2023).

6. K. McKee, H. Abdulla, L. O’Reilly, B. D. Walker, Cycling of labile and recalcitrant carboxyl-rich alicyclic molecules and carbohydrates in Baffin Bay. Nat Commun 15, 8735 (2024).

7. C. L. Follett, D. J. Repeta, D. H. Rothman, L. Xu, C. Santinelli, Hidden cycle of dissolved organic carbon in the deep ocean. Proceedings of the National Academy of Sciences 111, 16706–16711 (2014).

8. E. B. Kujawinski, The Impact of Microbial Metabolism on Marine Dissolved Organic Matter. Ann Rev Mar Sci 3, 567–599 (2011).

9. M. A. Moran, et al., Microbial metabolites in the marine carbon cycle. Nat Microbiol 7, 508–523 (2022).

10. E. B. Graham, J. E. Knelman, Implications of Soil Microbial Community Assembly for Ecosystem Restoration: Patterns, Process, and Potential. Microb Ecol 85, 809–819 (2023).

11. Q. Chen, et al., Correspondence between DOM molecules and microbial community in a subtropical coastal estuary on a spatiotemporal scale. Environ Int 154, 106558 (2021).

12. A. von Jackowski, et al., Seasonality of amino acid enantiomers and microbial communities at MOLA time series in the NW Mediterranean Sea. Org Geochem 196, 104839 (2024).

13. J. E. Goldford, et al., Emergent simplicity in microbial community assembly. Science 361, 469–474 (2018).

14. V. J. Coles, et al., Ocean biogeochemistry modeled with emergent trait-based genomics. Science 358, 1149–1154 (2017).

15. R. E. Danczak, et al., Ecological theory applied to environmental metabolomes reveals compositional divergence despite conserved molecular properties. Science of The Total Environment 788, 147409 (2021).

16. S. Louca, et al., Function and functional redundancy in microbial systems. Nat Ecol Evol 2, 936–943 (2018).

17. R. L. Bier, et al., Linking microbial community structure and microbial processes: an empirical and conceptual overview. FEMS Microbiol Ecol 91 (2015).

18. H. Fu, M. Uchimiya, J. Gore, M. A. Moran, Ecological drivers of bacterial community assembly in synthetic phycospheres. Proceedings of the National Academy of Sciences 117, 3656–3662 (2020).

19. A. D. Steen, et al., Analytical and Computational Advances, Opportunities, and Challenges in Marine Organic Biogeochemistry in an Era of “Omics.” Front Mar Sci 7, 718 (2020).

20. B. P. Durham, et al., Chemotaxonomic patterns in intracellular metabolites of marine microbial plankton. Front Mar Sci 9, 864796 (2022).

21. K. R. Heal, et al., Marine Community Metabolomes Carry Fingerprints of Phytoplankton Community Composition. mSystems 6 (2021).

22. E. B. Kujawinski, et al., Metabolite diversity among representatives of divergent *Prochlorococcus* ecotypes. mSystems 8 (2023).

23. S. A. Henson, et al., Uncertain response of ocean biological carbon export in a changing world. Nat Geosci 15, 248–254 (2022).

24. M. A. Moran, et al., The Ocean’s labile DOC supply chain. Limnol Oceanogr 67, 1007– 1021 (2022).

25. M. Huelsmann, M. Ackermann, Community instability in the microbial world. Science 378, 29–30 (2022).

26. A. F. Michaels, A. H. Knap, Overview of the U.S. JGOFS Bermuda Atlantic Time-series Study and the Hydrostation S program. Deep Sea Research Part II: Topical Studies in Oceanography 43, 157–198 (1996).

27. N. R. Bates, R. J. Johnson, Acceleration of ocean warming, salinification, deoxygenation and acidification in the surface subtropical North Atlantic Ocean. Commun Earth Environ 1, 1–12 (2020).

28. D. K. Steinberg, et al., Overview of the US JGOFS Bermuda Atlantic Time-series Study (BATS): A decade-scale look at ocean biology and biogeochemistry. Deep Sea Res II: Top Studies in Oceanography 48, 1405–1447 (2001).

29. M. W. Lomas, et al., Two decades and counting: 24-years of sustained open ocean biogeochemical measurements in the Sargasso Sea. Deep Sea Research Part II: Topical Studies in Oceanography 93, 16–32 (2013).

30. D. A. Hansell, C. A. Carlson, Biogeochemistry of total organic carbon and nitrogen in the Sargasso Sea: control by convective overturn. Deep Sea Research Part II: Topical Studies in Oceanography 48, 1649–1667 (2001).

31. S. Liu, et al., Linkages Among Dissolved Organic Matter Export, Dissolved Metabolites, and Associated Microbial Community Structure Response in the Northwestern Sargasso Sea on a Seasonal Scale. Front Microbiol 13, 407 (2022).

32. C. Carlson, H. Ducklow, A. Michaels, Annual flux of dissolved organic carbon from the euphotic zone in the northwestern Sargasso Sea. Nature Letters 371, 405–408 (1994).

33. S. J. Giovannoni, K. L. Vergin, Seasonality in ocean microbial communities. Science 335, 671–676 (2012).

34. A. Roesch, H. Schmidbauer, WaveletComp: Computational Wavelet Analysis. R package version 1.1. [Preprint] (2018). Available at: https://cran.r-project.org/package=WaveletComp [Accessed 5 December 2022].

35. J. Merder, et al., Dissolved organic compounds with synchronous dynamics share chemical properties and origin. Limnol Oceanogr 66, 4001–4016 (2021).

36. R. M. Boiteau, et al., Relating Molecular Properties to the Persistence of Marine Dissolved Organic Matter with Liquid Chromatography-Ultrahigh-Resolution Mass Spectrometry. Environ Sci Technol 58, 3267–3277 (2023).

37. A. M. Martin-Platero, et al., High resolution time-series reveals cohesive but short-lived communities in coastal plankton. Nat Commun 9, 1–11 (2018).

38. L. M. Bolaños, et al., Influence of short and long term processes on SAR11 communities in open ocean and coastal systems. ISME Communications 2, 1–11 (2022).

39. D. Broadhurst, et al., Guidelines and considerations for the use of system suitability and quality control samples in mass spectrometry assays applied in untargeted clinical metabolomic studies. Metabolomics 14, 1–17 (2018).

40. S. Roshan, T. DeVries, Efficient dissolved organic carbon production and export in the oligotrophic ocean. Nat Commun 8, 2036 (2017).

41. S. Sunagawa, et al., Structure and function of the global ocean microbiome. Science 348 (2015).

42. M. Gonsior, et al., Optical properties and molecular differences in dissolved organic matter at the Bermuda Atlantic and Hawai’i ALOHA time-series stations. Environmental Science: Advances 3, 717–731 (2024).

43. J. A. Ivory, D. K. Steinberg, R. J. Latour, Diel, seasonal, and interannual patterns in mesozooplankton abundance in the Sargasso Sea. ICES Journal of Marine Science 76, 217– 231 (2019).

44. M. R. Landry, H. Al-Mutairi, K. E. Selph, S. Christensen, S. Nunnery, Seasonal patterns of mesozooplankton abundance and biomass at Station ALOHA. Deep Sea Research Part II: Topical Studies in Oceanography 48, 2037–2061 (2001).

45. J. H. W. Saw, et al., Pangenomics analysis reveals diversification of enzyme families and niche specialization in globally abundant SAR202 bacteria. mBio 11 (2020).

46. D. K. Steinberg, et al., Zooplankton vertical migration and the active transport of dissolved organic and inorganic carbon in the Sargasso Sea. Deep Sea Research Part I: Oceanographic Research Papers 47, 137–158 (2000).

47. A. E. Maas, et al., Migratory Zooplankton Excreta and Its Influence on Prokaryotic Communities. Front Mar Sci 7, 573268 (2020).

48. I. Koester, et al., Illuminating the dark metabolome of Pseudo-nitzschia–microbiome associations. Environ Microbiol 24, 5408–5424 (2022).

49. L. W. Sumner, et al., Proposed minimum reporting standards for chemical analysis: Chemical Analysis Working Group (CAWG) Metabolomics Standards Initiative (MSI). Metabolomics 3, 211–221 (2007).

50. B. Gebser, K. Thume, M. Steinke, G. Pohnert, Phytoplankton-derived zwitterionic gonyol and dimethylsulfonioacetate interfere with microbial dimethylsulfoniopropionate sulfur cycling. Microbiologyopen 9, e1014 (2020).

51. W. M. Johnson, M. C. Kido Soule, E. B. Kujawinski, Extraction efficiency and quantification of dissolved metabolites in targeted marine metabolomics. Limnol. Oceanogr.: Methods 15, 417–428 (2017).

52. B. Widner, M. C. Kido Soule, F. X. Ferrer-González, M. A. Moran, E. B. Kujawinski, Quantification of Amine- And Alcohol-Containing Metabolites in Saline Samples Using Pre-extraction Benzoyl Chloride Derivatization and Ultrahigh Performance Liquid Chromatography Tandem Mass Spectrometry (UHPLC MS/MS). Anal Chem 93, 4809– 4817 (2021).

53. J. S. Sacks, K. R. Heal, A. K. Boysen, L. T. Carlson, A. E. Ingalls, Quantification of dissolved metabolites in environmental samples through cation-exchange solid-phase extraction paired with liquid chromatography–mass spectrometry. Limnol Oceanogr Methods 20, 683–700 (2022).

54. T. Dittmar, B. Koch, N. Hertkorn, G. Kattner, A simple and efficient method for the solid-phase extraction of dissolved organic matter (SPE-DOM) from seawater. Limnol Oceanogr Methods 6, 230–235 (2008).

55. S. K. Bercovici, M. C. Arroyo, D. De Corte, T. Yokokawa, D. A. Hansell, Limited utilization of extracted dissolved organic matter by prokaryotic communities from the subtropical North Atlantic. Limnol Oceanogr 66, 2509–2520 (2021).

56. S. Liu, et al., Stable Isotope Probing Identifies Bacterioplankton Lineages Capable of Utilizing Dissolved Organic Matter Across a Range of Bioavailability. Front Microbiol 11, 2364 (2020).

57. E. Jerusalén-Lleó, M. Nieto-Cid, I. Fuentes-Santos, T. Dittmar, X. A. Álvarez-Salgado, Solid phase extraction of ocean dissolved organic matter with PPL cartridges: efficiency and selectivity. Front Mar Sci 10, 1159762 (2023).

58. K. Longnecker, et al., Seasonal and daily patterns in known dissolved metabolites in the northwestern Sargasso Sea. Limnol Oceanogr 69, 449–466 (2024).

59. C. L. Fiore, K. Longnecker, M. C. Kido Soule, E. B. Kujawinski, Release of Ecologically Relevant Metabolites by the Cyanobacterium, *Synechococcus elongatus* CCMP 1631. Environ Microbiol 17 (2015).

60. L. Weber, et al., Extracellular Reef Metabolites Across the Protected Jardines de la Reina, Cuba Reef System. Front Mar Sci 7, 1063 (2020).

61. A. Vorobev, et al., Identifying labile DOM components in a coastal ocean through depleted bacterial transcripts and chemical signals. Environ Microbiol 20, 1–19 (2018).

62. D. A. Hansell, Recalcitrant dissolved organic carbon fractions. Ann Rev Mar Sci 5, 421– 445 (2013).

63. N. M. Levine, et al., Revising upper-ocean sulfur dynamics near Bermuda: New lessons from 3 years of concentration and rate measurements. Environmental Chemistry 13, 302– 313 (2015).

64. R. Kiene, L. Linn, The fate of dissolved dimethylsulfoniopropionate (DMSP) in seawater: Tracer studies using 35 S-DMSP. Geochim Cosmochim Acta 64, 2797–2810 (2000).

65. E. F. DeLong, et al., Community genomics among stratified microbial assemblages in the ocean’s interior. Science 311, 496–503 (2006).

66. R. R. Malmstrom, et al., Temporal dynamics of *Prochlorococcus* ecotypes in the Atlantic and Pacific oceans. ISME J 4, 1252–1264 (2010).

67. S. J. Giovannoni, M. S. Rappé, K. L. Vergin, N. L. Adair, 16S rRNA genes reveal stratified open ocean bacterioplankton populations related to the Green Non-Sulfur bacteria. Proceedings of the National Academy of Sciences 93, 7979–7984 (1996).

68. J. W. Becker, et al., Closely related phytoplankton species produce similar suites of dissolved organic matter. Front Microbiol 5 (2014).

69. K. L. Vergin, B. Done, C. A. Carlson, S. J. Giovannoni, Spatiotemporal distributions of rare bacterioplankton populations indicate adaptive strategies in the oligotrophic ocean. Aquatic Microbial Ecology 71, 1–13 (2013).

70. A. A. Larkin, et al., Subtle biogeochemical regimes in the Indian Ocean revealed by spatial and diel frequency of Prochlorococcus haplotypes. Limnol Oceanogr 65, S220–S232 (2020).

71. C. A. Lozupone, J. I. Stombaugh, J. I. Gordon, J. K. Jansson, R. Knight, Diversity, stability and resilience of the human gut microbiota. Nature 489, 220–230 (2012).

72. S. Lambert, J.-C. Lozano, F.-Y. Bouget, P. E. Galand, Seasonal marine microorganisms change neighbours under contrasting environmental conditions. Environmental Microbiology 23, 2592-2604 (2021).

73. C. S. Ward, et al., Annual community patterns are driven by seasonal switching between closely related marine bacteria. ISME J 11, 1412–1422 (2017).

74. B. Segerman, The genetic integrity of bacterial species: the core genome and the accessory genome, two different stories. Front Cell Infect Microbiol 2, 116 (2012).

75. C. Burke, P. Steinberg, D. Rusch, S. Kjelleberg, T. Thomas, Bacterial community assembly based on functional genes rather than species. Proceedings of the National Academy of Sciences 108, 14288–14293 (2011).

76. S. D. Allison, J. B. H. Martiny, Resistance, resilience, and redundancy in microbial communities. Proceedings of the National Academy of Sciences 105, 11512–11519 (2008).

77. S. Louca, P. L, M. Doebeli, Decoupling function and taxonomy in the global ocean microbiome. Science 353, 1272–1277 (2016).

78. E. L. McParland, H. Alexander, W. M. Johnson, The Osmolyte Ties That Bind: Genomic Insights Into Synthesis and Breakdown of Organic Osmolytes in Marine Microbes. Front Mar Sci 8, 732 (2021).

79. T. M. Royalty, A. D. Steen, Contribution Evenness: A functional redundancy metric sensitive to trait stability in microbial communities. bioRxiv (2021). 10.1101/2020.04.22.054593.

80. A. H. Treusch, et al., Seasonality and vertical structure of microbial communities in an ocean gyre. ISME J 3, 1148–1163 (2009).

81. C. A. Carlson, et al., Seasonal dynamics of SAR11 populations in the euphotic and mesopelagic zones of the northwestern Sargasso Sea. ISME J 3, 283–295 (2009).

82. K. L. Vergin, et al., High-resolution SAR11 ecotype dynamics at the Bermuda Atlantic Time-series Study site by phylogenetic placement of pyrosequences. ISME J 7, 1322–1332 (2013).

83. W. H. Cheng, C. H. Hsieh, C. W. Chang, F. K. Shiah, T. Miki, New index of functional specificity to predict the redundancy of ecosystem functions in microbial communities. FEMS Microbiol Ecol 98 (2022).

84. N. R. Bates, R. J. Johnson, Acceleration of ocean warming, salinification, deoxygenation and acidification in the surface subtropical North Atlantic Ocean. Commun Earth Environ 1, 1–12 (2020).

85. S. P. Hubbell, The Unified Neutral Theory of Biodiversity and Biogeography (Princeton University Press, 2001).

86. N. M. Levine, M. A. Doblin, S. Collins, Reframing trait trade-offs in marine microbes. Commun Earth Environ 5, 1–6 (2024).

87. B. B. Cael, S. Dutkiewicz, S. Henson, Abrupt shifts in 21st-century plankton communities. Sci Adv 7, 8593–8622 (2021).

88. P. W. Boyd, R. Strzepek, F. Fu, D. a. Hutchins, Environmental control of open-ocean phytoplankton groups: Now and in the future. Limnol Oceanogr 55, 1353–1376 (2010).

89. S. Dutkiewicz, J. R. Scott, M. J. Follows, Winners and losers: Ecological and biogeochemical changes in a warming ocean. Global Biogeochem Cycles 27, 463–477 (2013).

90. P. Flombaum, et al., Present and future global distributions of the marine Cyanobacteria *Prochlorococcus* and *Synechococcus*. Proceedings of the National Academy of Sciences 110, 9824–9829 (2013).

91. M. C. Kido Soule, K. Longnecker, W. M. Johnson, E. B. Kujawinski, Environmental metabolomics: Analytical strategies. Mar Chem 177, 374–387 (2015).

92. K. Longnecker, Dissolved organic matter in newly formed sea ice and surface seawater. Geochim Cosmochim Acta 171, 39–49 (2015).

93. M. C. Chambers, et al., A cross-platform toolkit for mass spectrometry and proteomics. Nat Biotechnol 30, 918–920 (2012).

94. C. A. Smith, E. J. Want, G. O’Maille, R. Abagyan, G. Siuzdak, XCMS: Processing mass spectrometry data for metabolite profiling using nonlinear peak alignment, matching, and identification. Anal Chem 78, 779–787 (2006).

95. C. Kuhl, R. Tautenhahn, C. Böttcher, T. R. Larson, S. Neumann, CAMERA: An integrated strategy for compound spectra extraction and annotation of LC/MS data sets. Anal Chem 84, 283 (2012).

96. R. Wehrens, et al., Improved batch correction in untargeted MS-based metabolomics. Metabolomics 12 (2016).

97. W. B. Dunn, et al., Procedures for large-scale metabolic profiling of serum and plasma using gas chromatography and liquid chromatography coupled to mass spectrometry. Nat Protoc 6, 1060–1083 (2011).

98. L. F. Nothias, et al., Feature-based molecular networking in the GNPS analysis environment. Nat Methods 17, 905–908 (2020).

99. B. J. Callahan, P. J. McMurdie, S. P. Holmes, Exact sequence variants should replace operational taxonomic units in marker-gene data analysis. ISME J 11, 2639–2643 (2017).

100. C. Quast, et al., The SILVA ribosomal RNA gene database project: improved data processing and web-based tools. Nucleic Acids Res 41 (2013).

101. P. J. McMurdie, S. Holmes, phyloseq: An R Package for Reproducible Interactive Analysis and Graphics of Microbiome Census Data. PLoS One 8, e61217 (2013).

102. T. Aramaki, et al., KofamKOALA: KEGG Ortholog assignment based on profile HMM and adaptive score threshold. Bioinformatics 36, 2251–2252 (2020).

103. M. Kanehisa, S. Goto, KEGG: Kyoto Encyclopedia of Genes and Genomes. Nucleic Acids Res 28, 27–30 (2000).

104. J. Vollmers, S. Wiegand, F. Lenk, A. K. Kaster, How clear is our current view on microbial dark matter? (Re-)assessing public MAG & SAG datasets with MDMcleaner. Nucleic Acids Res 50, e76–e76 (2022).

105. S. J. Biller, et al., Data Descriptor : Marine microbial metagenomes sampled across space and time. Sci Data 5, 180176 (2018).

## References

1. M. C. Chambers, et al., A cross-platform toolkit for mass spectrometry and proteomics. Nat Biotechnol 30, 918–920 (2012).

2. W. B. Dunn, et al., Procedures for large-scale metabolic profiling of serum and plasma using gas chromatography and liquid chromatography coupled to mass spectrometry. Nat Protoc 6, 1060–1083 (2011).

3. J. T. Prince, E. M. Marcotte, Chromatographic alignment of ESI-LC-MS proteomics data sets by ordered bijective interpolated warping. Anal Chem 78, 6140–6152 (2006).

4. C. A. Smith, E. J. Want, G. O’Maille, R. Abagyan, G. Siuzdak, XCMS: Processing mass spectrometry data for metabolite profiling using nonlinear peak alignment, matching, and identification. Anal Chem 78, 779–787 (2006).

5. C. Kuhl, R. Tautenhahn, C. Böttcher, T. R. Larson, S. Neumann, CAMERA: An integrated strategy for compound spectra extraction and annotation of LC/MS data sets. Anal Chem 84, 283 (2012).

6. C. Schiffman, et al., Filtering procedures for untargeted LC-MS metabolomics data. BMC Bioinformatics 20, 1–10 (2019).

7. M. Zark, J. Christoffers, T. Dittmar, Molecular properties of deep-sea dissolved organic matter are predictable by the central limit theorem: Evidence from tandem FT-ICR-MS. Mar Chem 191, 9–15 (2017).

8. K. Lu, X. Li, H. Chen, Z. Liu, Constraints on isomers of dissolved organic matter in aquatic environments: Insights from ion mobility mass spectrometry. Geochim Cosmochim Acta 308, 353–372 (2021).

9. L. W. Sumner, et al., Proposed minimum reporting standards for chemical analysis: Chemical Analysis Working Group (CAWG) Metabolomics Standards Initiative (MSI). Metabolomics 3, 211–221 (2007).

10. D. Broadhurst, et al., Guidelines and considerations for the use of system suitability and quality control samples in mass spectrometry assays applied in untargeted clinical metabolomic studies. Metabolomics 14, 1–17 (2018).

11. Z. Zhang, et al., Reducing Quantitative Uncertainty Caused by Data Processing in Untargeted Metabolomics. Anal Chem 96, 3727–3732 (2024).

12. R. D. Beger, et al., Towards quality assurance and quality control in untargeted metabolomics studies. Metabolomics 15, 1–5 (2019).

13. P. Stincone, et al., Evaluation of Data-Dependent MS/MS Acquisition Parameters for Non-Targeted Metabolomics and Molecular Networking of Environmental Samples: Focus on the Q Exactive Platform. Anal Chem 95, 12673–12682 (2023).

14. A. Spielmeyer, G. Pohnert, Direct quantification of dimethylsulfoniopropionate (DMSP) with hydrophilic interaction liquid chromatography/mass spectrometry. J Chromatogr B Analyt Technol Biomed Life Sci 878, 3238–3242 (2010).

15. B. Gebser, G. Pohnert, Synchronized regulation of different zwitterionic metabolites in the osmoadaption of phytoplankton. Mar Drugs 11, 2168–2182 (2013).

16. B. Gebser, K. Thume, M. Steinke, G. Pohnert, Phytoplankton-derived zwitterionic gonyol and dimethylsulfonioacetate interfere with microbial dimethylsulfoniopropionate sulfur cycling. Microbiologyopen 9, e1014 (2020).

17. K. R. Heal, et al., Marine Community Metabolomes Carry Fingerprints of Phytoplankton Community Composition. mSystems 6 (2021).

18. J. S. Sacks, K. R. Heal, A. K. Boysen, L. T. Carlson, A. E. Ingalls, Quantification of dissolved metabolites in environmental samples through cation-exchange solid-phase extraction paired with liquid chromatography–mass spectrometry. Limnol Oceanogr Methods 20, 683–700 (2022).

19. C. Arnosti, et al., The Biogeochemistry of Marine Polysaccharides: Sources, Inventories, and Bacterial Drivers of the Carbohydrate Cycle. Ann Rev Mar Sci 13, 81–8 (2021).

20. M. Wang, et al., The great Atlantic Sargassum belt. Science (1979) 364, 83–87 (2019).

21. W. Helbert, Marine polysaccharide sulfatases. Front Mar Sci 4, 6 (2017).

22. A. G. Hettle, C. J. Vickers, A. B. Boraston, Sulfatases: Critical Enzymes for Algal Polysaccharide Processing. Front Plant Sci 13 (2022).

23. J. H. W. Saw, et al., Pangenomics analysis reveals diversification of enzyme families and niche specialization in globally abundant SAR202 bacteria. mBio 11 (2020).

24. B. A. S. Van Mooy, G. Rocap, H. F. Fredricks, C. T. Evans, A. H. Devol, Sulfolipids dramatically decrease phosphorus demand by picocyanobacteria in oligotrophic marine environments. Proceedings of the National Academy of Sciences 103, 8607–12 (2006).

25. J. C. Argüelles, Physiological roles of trehalose in bacteria and yeasts: a comparative analysis. Arch Microbiol 174, 217–224 (2000).

26. N. Pade, J. Compaoré, S. Klähn, L. J. Stal, M. Hagemann, The marine cyanobacterium *Crocosphaera watsonii* WH8501 synthesizes the compatible solute trehalose by a laterally acquired OtsAB fusion protein. Environ Microbiol 14, 1261–1271 (2012).

27. C. McLean, et al., Harmful Algal Bloom-Forming Organism Responds to Nutrient Stress Distinctly From Model Phytoplankton. bioRxiv 2021.02.08.430350 (2021). 10.1101/2021.02.08.430350.

28. A. K. Boysen, et al., Particulate Metabolites and Transcripts Reflect Diel Oscillations of Microbial Activity in the Surface Ocean. mSystems 6 (2021).

29. O. Levitan, et al., Remodeling of intermediate metabolism in the diatom *Phaeodactylum tricornutum* under nitrogen stress. Proceedings of the National Academy of Sciences 112 (2015).

30. S. E. Noell, et al., SAR11 cells rely on enzyme multifunctionality to metabolize a range of polyamine compounds. mBio 12 (2021).

31. C. Koedooder, et al., The role of the glyoxylate shunt in the acclimation to iron limitation in marine heterotrophic bacteria. Front Mar Sci 5, 435 (2018).

