## Supplemental Material for "Seasonal patterns of exometabolites depend on microbial functions in the oligotrophic ocean"

Table S2: List of metabolic KOs queried in all surface metagenomes for trehalose and succinate production or consumption.

Table S3: Metagenome assemblies queried for functional redundancy analyses.

Table S4: A mix of stable isotope labeled internal standards was added to every sample of the exometabolome. The label reflects which element was isotopically heavy. Ng/ml reflects the concentration added to each sample. Ion mode reflects which ionization mode the standard was detected in. The retention time reflects where the standard was detected in the chromatogram. The exact mass was calculated based on monoisotopic element composition. All standards were detected as either [M+H] or [M-H] adducts.

Fig S1

A original time series

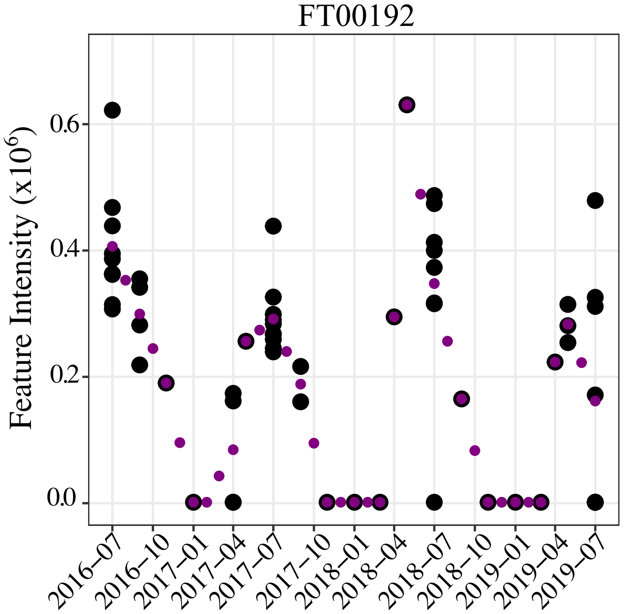


B wavelet transformation of time series

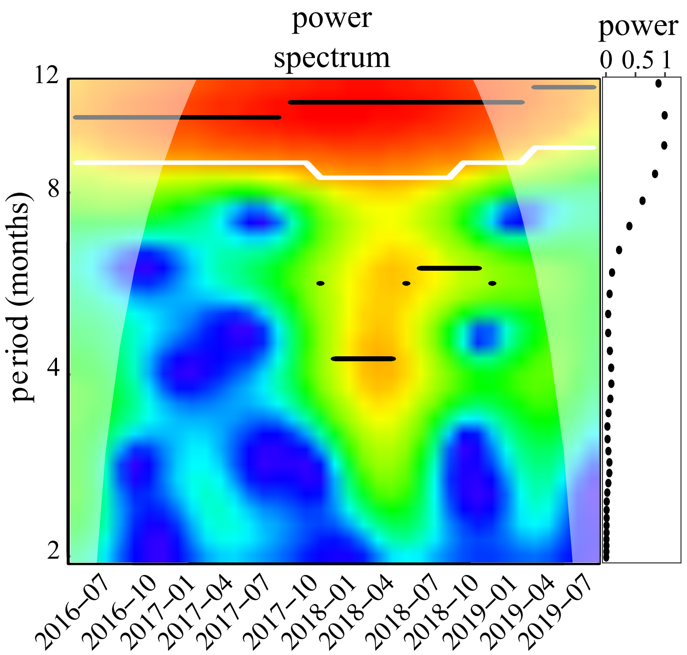


C wavelet reconstructions were used to predict seasonal maxima

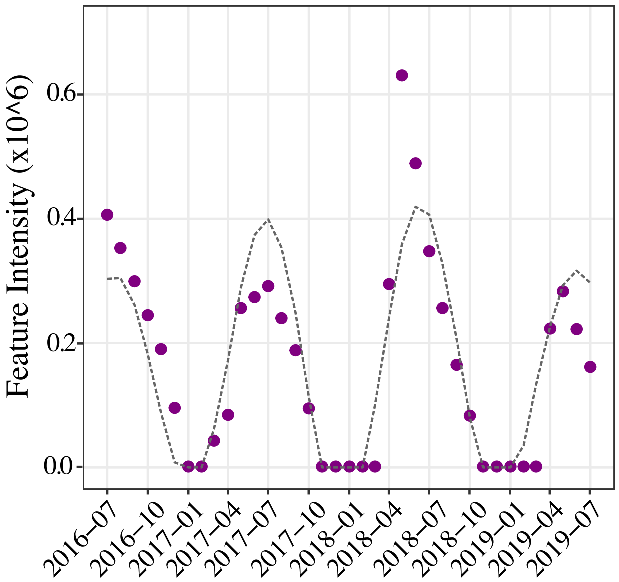

Fig S2

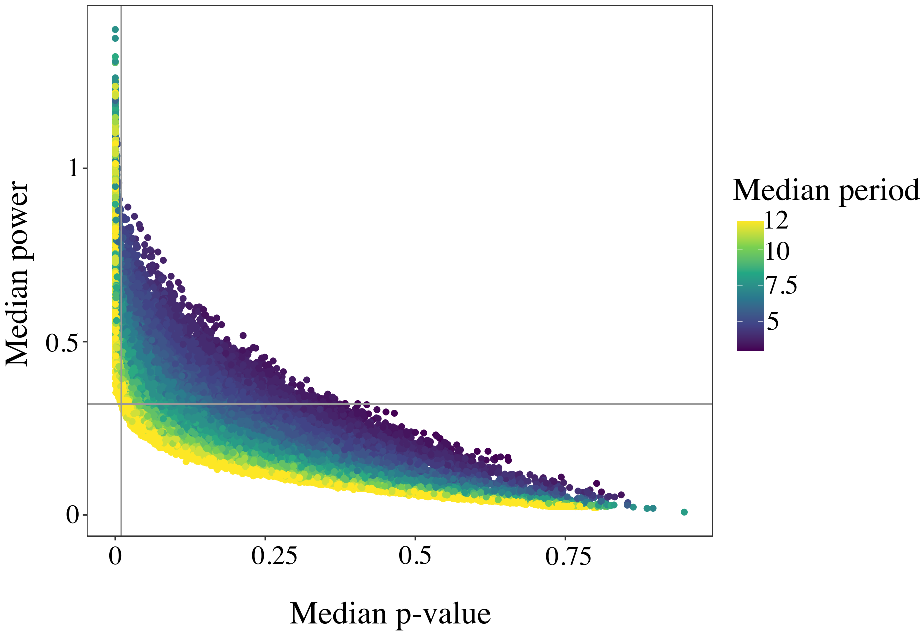

Median power
1
0.5
0
0
0.25
0.5
0.75
Median p-value
Median period
12
10
7.5
5

Fig S3

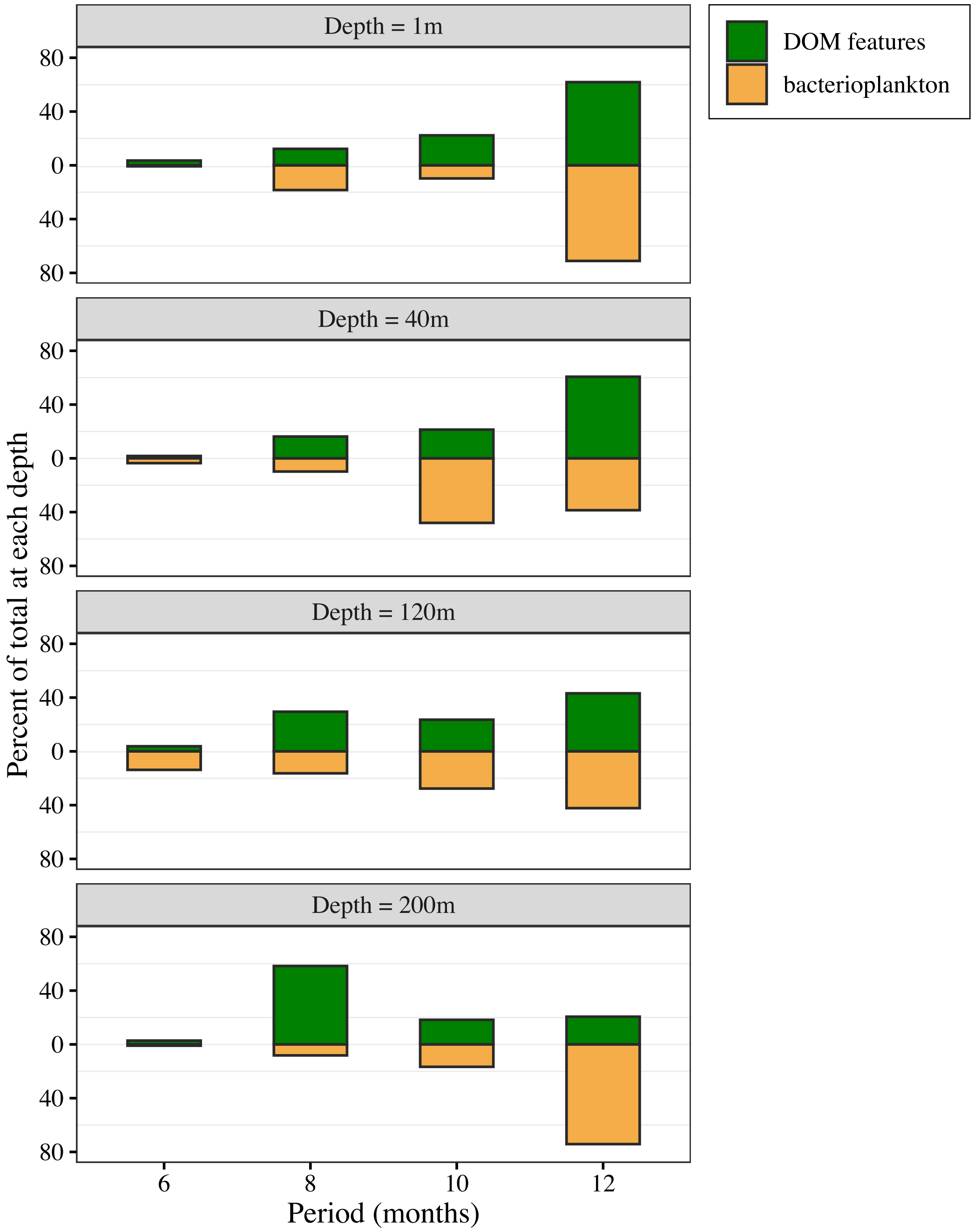

Depth = 1m
Depth = 40m
Depth = 120m
Depth = 200m
80
40
0
40
80
Percent of total at each depth
6
8
10
12
Period (months)
DOM features
bacterioplankton

Fig S4

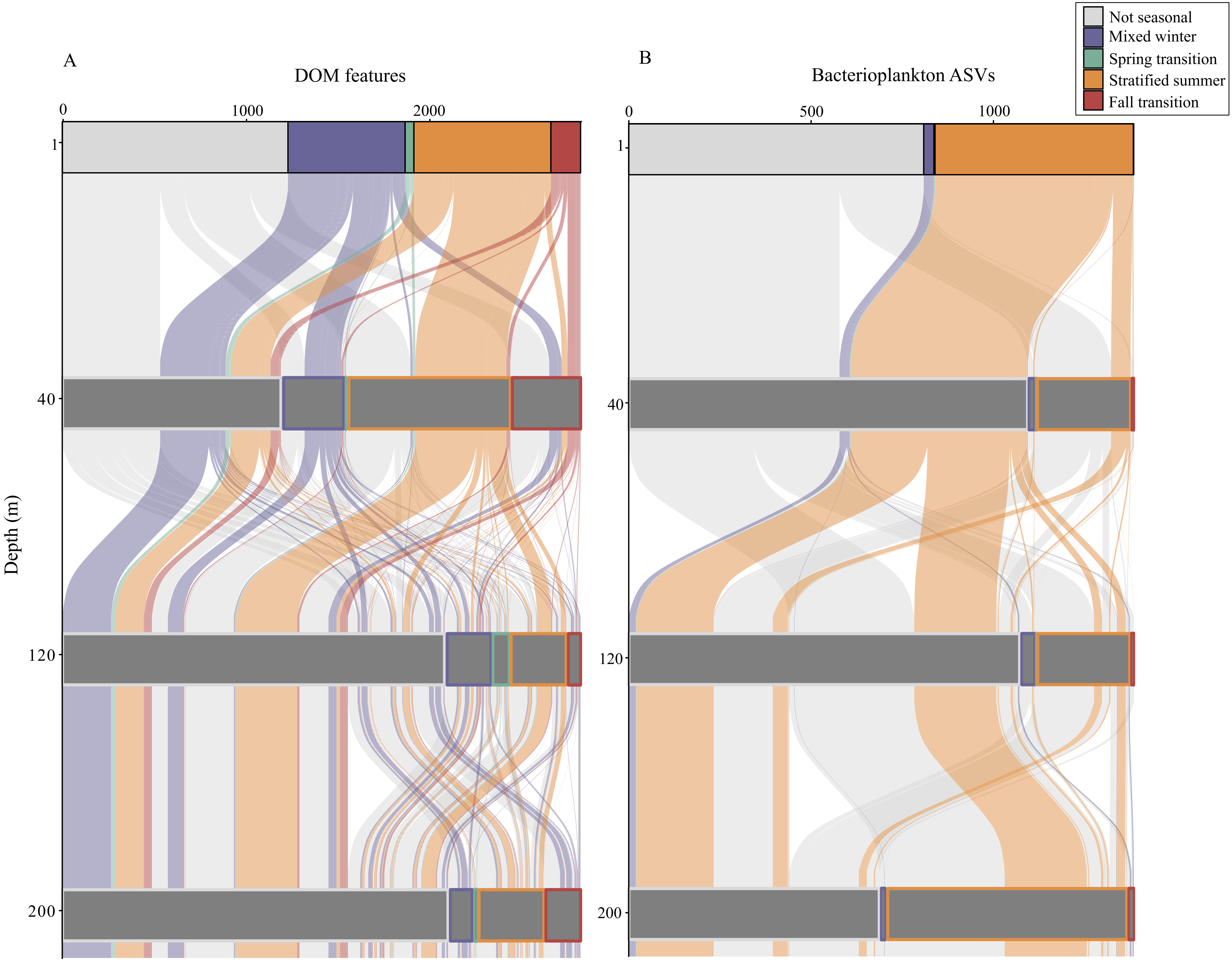

Not seasonal
Mixed winter
Spring transition
Stratified summer
Fall transition
A
DOM features
0
1000
2000
B
Bacterioplankton ASVs
0
500
1000
1
40
120
200
Depth (m)

SFig 5

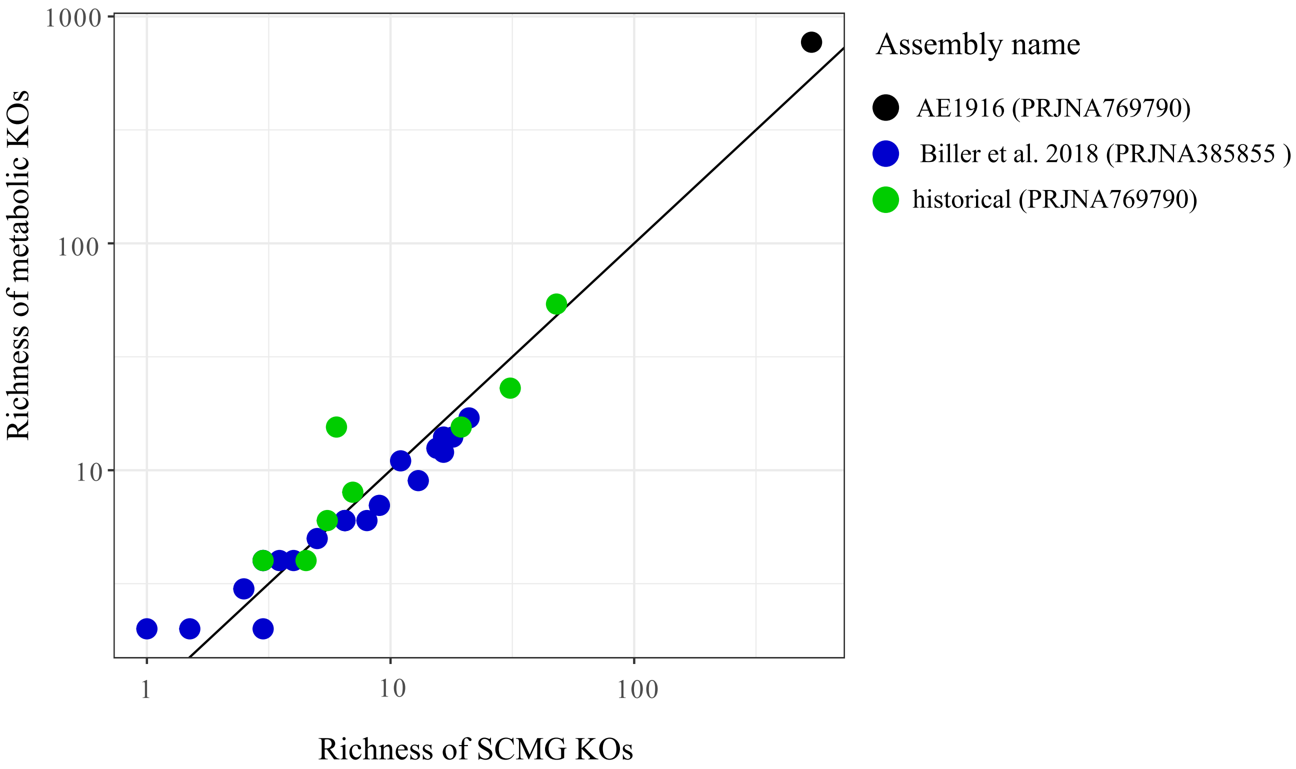
1000
100
10
1
10
100
Richness of metabolic KOs
Richness of SCMG KOs
Assembly name
AE1916 (PRJNA769790)
Biller et al. 2018 (PRJNA385855 )
historical (PRJNA769790)

succinate



A MS2 spectra comparison between sample (black) and reference (green)

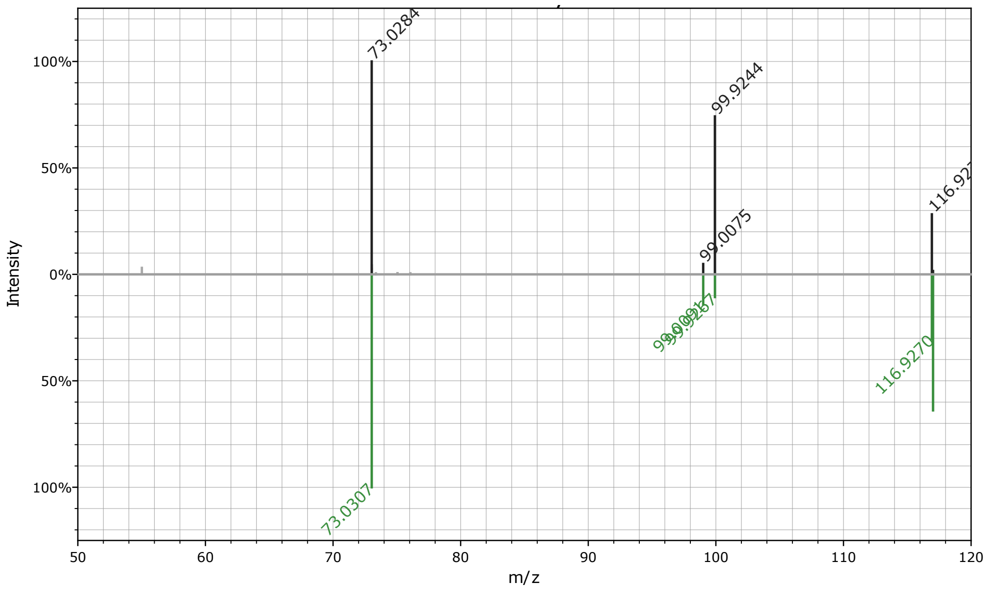


B EIC of samples (blue) and succinate standard (black)

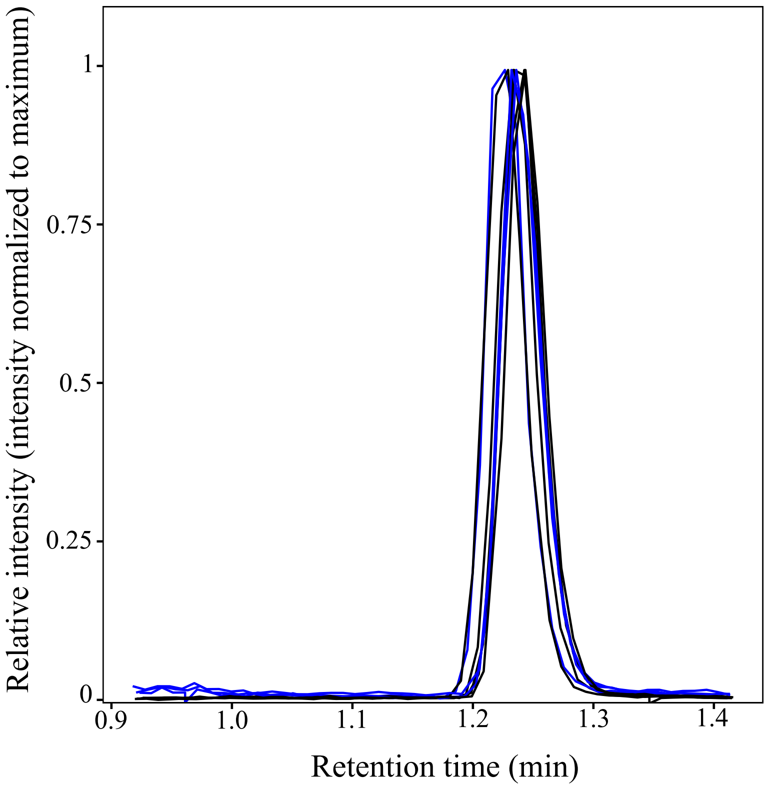

trehalose

Fig S7

A MS2 spectra comparison between sample (black) and reference (green)

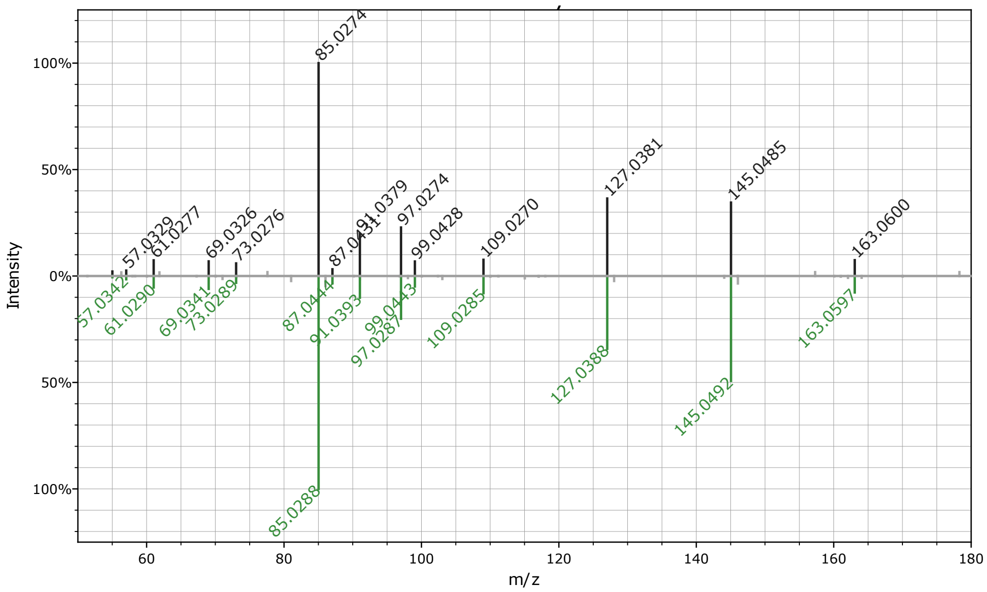


B EIC of samples (blue) and trehalose and sucrose standard (black)

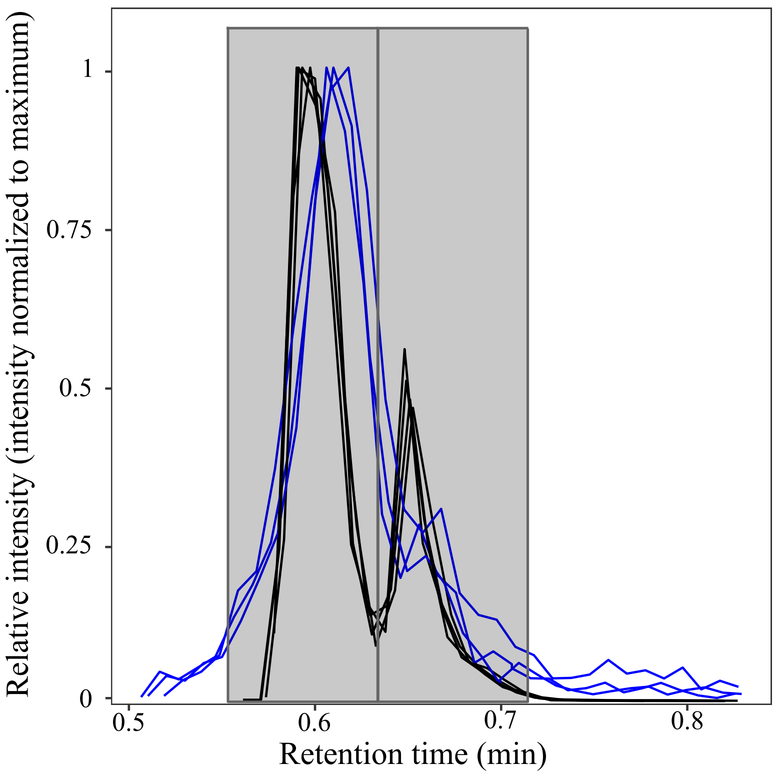

gonyol



A MS2 spectra comparison between sample (black) and reference (green)

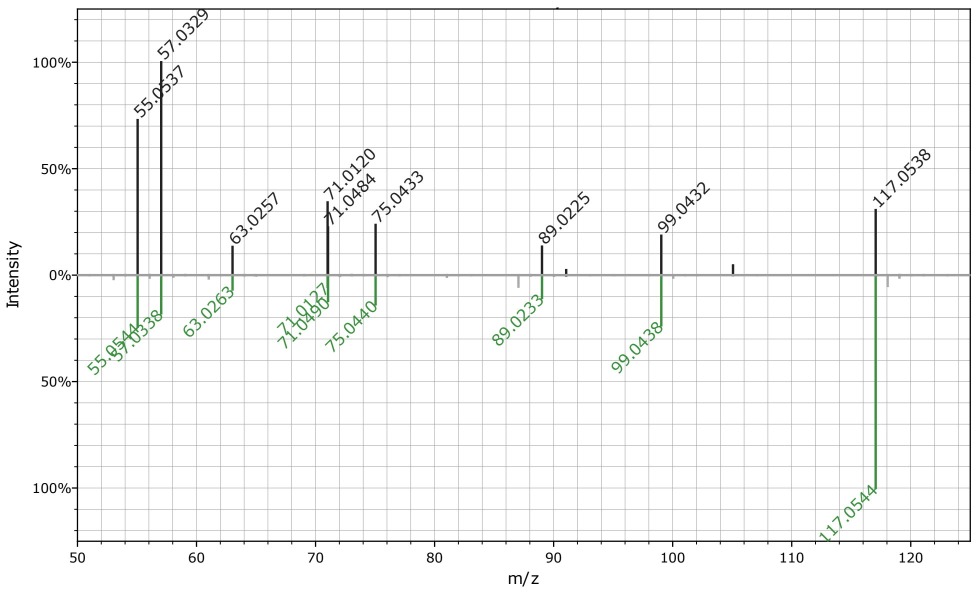


B EIC of sample (blue) and gonyol standard (black)

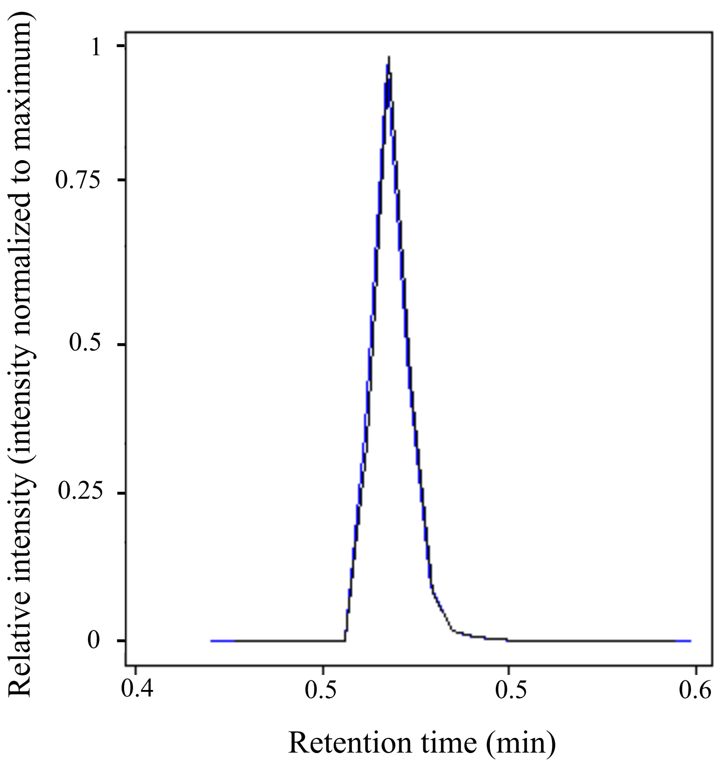

glucose 6-sulfate



A MS2 spectra comparison between sample (black) and reference (green)

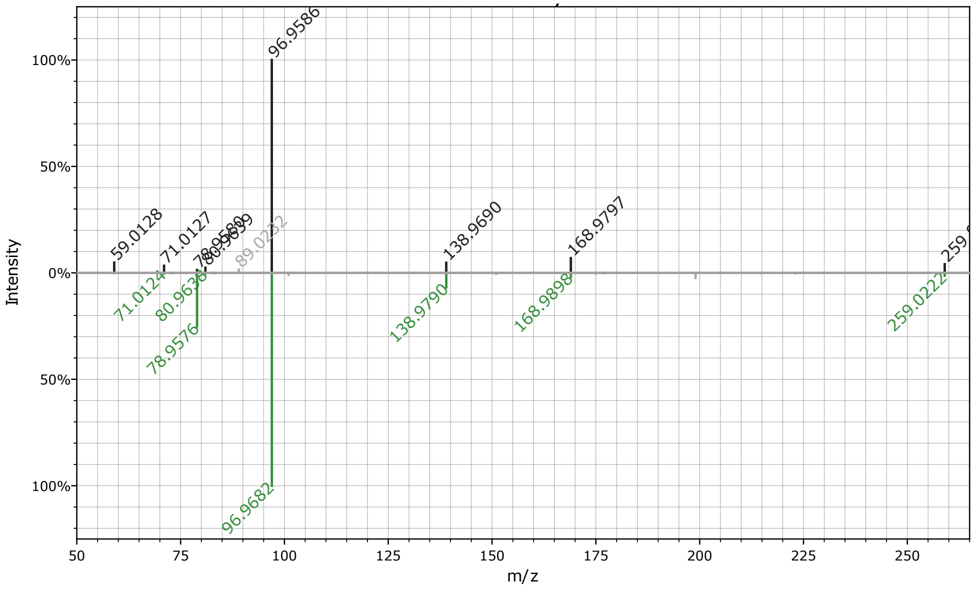

| Compound | Formula | Id level | Ion mode | m/z | Adduct | Ret time (min) |
|---|---|---|---|---|---|---|
| gonyol | $C_7H_{14}O_3S$ | 1 | Pos | 179.0736 | $[M+H]^+$ | 0.57 |
| trehalose | $C_{12}H_{22}O_{11}$ | 1 | Pos | 360.1504 | $[M+NH_4]^+$ | 0.61 |
| succinate | $C_4H_6O_4$ | 1 | Neg | 117.0192 | $[M-H]^-$ | 1.24 |
| glucose 6-sulfate | $C_6H_{12}O_9S$ | 2 | Neg | 259.0128 | $[M-H]^-$ | 0.61 |

| compound_name | KO | enzyme_name | direction | group |
|---|---|---|---|---|
| Succinate | K00135 | succinate-semialdehyde dehydrogenase / glutarate-semialdehyde dehydrogenase [EC:1.2.1.16 1.2.1.79 1.2.1.20] | production | succinate-semialdehyde dehydrogenase |
| Succinate | K00139 | succinate-semialdehyde dehydrogenase [EC:1.2.1.24] | production | succinate-semialdehyde dehydrogenase |
| Succinate | K17761 | succinate-semialdehyde dehydrogenase, mitochondrial [EC:1.2.1.24] | production | succinate-semialdehyde dehydrogenase |
| Succinate | K08324 | succinate-semialdehyde dehydrogenase [EC:1.2.1.16 1.2.1.24] | production | succinate-semialdehyde dehydrogenase |
| Succinate | K01902 | succinyl-CoA synthetase alpha subunit [EC:6.2.1.5] | production | succinyl CoA synthetase |
| Succinate | K01899 | succinyl-CoA synthetase alpha subunit [EC:6.2.1.4 6.2.1.5] | production | succinyl CoA synthetase |
| Succinate | K15737 | glutarate dioxygenase [EC:1.14.11.64] | production | glutarate oxidoreductase |
| Succinate | K18118 | succinyl-CoA:acetate CoA-transferase [EC:2.8.3.18] | production | succinyl CoA - acetate CoA transferase |
| Succinate | K00244 | succinate dehydrogenase flavoprotein subunit [EC:1.3.5.1] | consumption | succinate dehydrogenase |
| Succinate | K00234 | succinate dehydrogenase (ubiquinone) flavoprotein subunit [EC:1.3.5.1] | consumption | succinate dehydrogenase |
| Succinate | K00239 | succinate dehydrogenase flavoprotein subunit [EC:1.3.5.1] | consumption | succinate dehydrogenase |
| Trehalose | K13057 | trehalose synthase | production | glycosyltransferase |
| Trehalose | K05343 | maltose alpha-D-glucosyltransferase / alpha-amylase | production | glucosyltransferase |
| Trehalose | K01236 | maltooligosyltrehalose trehalohydrolase | production | trehalohydrolase |
| Trehalose | K01087 | trehalose 6-phosphate phosphatase | production | phosphatase |
| Trehalose | K22934 | alpha,alpha-trehalase | consumption | trehalase |
| Trehalose | K01194 | alpha,alpha-trehalase | consumption | trehalase |
| Trehalose | K05342 | alpha,alpha-trehalose phosphorylase | consumption | trehalose phosphorylase |

| | Assembly name | Date (year-month-day) | Depth (m) | sample |
|---|---|---|---|---|
| 1 | hist | 1997-09-01 | 1 | 108_0 |
| 2 | hist | 1998-02-01 | 1 | 113_0 |
| 3 | hist | 1999-11-01 | 1 | 134_0 |
| 4 | hist | 2000-01-01 | 1 | 136_0 |
| 5 | hist | 2000-03-01 | 1 | 138_0 |
| 6 | hist | 2001-08-01 | 1 | 155_0 |
| 7 | hist | 2002-05-01 | 1 | 164_0 |
| 8 | Biller | 2003-02-21 | 1 | SRR5720233 |
| 9 | Biller | 2003-03-22 | 1 | SRR5720238 |
| 10 | Biller | 2003-04-22 | 1 | SRR5720327 |
| 11 | Biller | 2003-05-20 | 1 | SRR5720283 |
| 12 | Biller | 2003-07-15 | 1 | SRR5720235 |
| 13 | Biller | 2003-08-12 | 1 | SRR5720286 |
| 14 | Biller | 2003-10-07 | 1 | SRR5720332 |
| 15 | Biller | 2003-11-04 | 1 | SRR5720276 |
| 16 | Biller | 2003-12-02 | 1 | SRR5720262 |
| 17 | hist | 2003-03-01 | 1 | 174A_0 |
| 18 | Biller | 2004-01-27 | 1 | SRR5720338 |
| 19 | Biller | 2004-02-24 | 1 | SRR5720322 |
| 20 | Biller | 2004-03-23 | 1 | SRR5720337 |
| 21 | Biller | 2004-04-21 | 1 | SRR5720256 |
| 22 | Biller | 2004-05-18 | 1 | SRR5720257 |
| 23 | Biller | 2004-06-15 | 1 | SRR5720260 |
| 24 | Biller | 2004-08-17 | 1 | SRR5720321 |
| 25 | Biller | 2004-09-14 | 1 | SRR5720251 |
| 26 | Biller | 2004-10-13 | 1 | SRR5720307 |
| 27 | Biller | 2004-11-12 | 1 | SRR5720278 |
| 28 | Biller | 2004-12-08 | 1 | SRR5720342 |
| 29 | Biller | 2009-07-14 | 1 | SRR6507279 |
| 30 | AE1916 | 2019-07-09 | 1 | 5_1_S27 |

| | | Positive ionization mode | | | |
|---|---|---|---|---|---|
| | | Detection | | RSD | |
| compound | label | blanks (n = 15) | samples (n = 256) | blanks | samples |
| leucine | D3 | 14 | 256 | 0.28 | 0.03 |
| methionine | D3 | 15 | 256 | 0.07 | 0.06 |
| phenyalanine | D8 | 15 | 256 | 0.03 | 0.07 |
| proline | 13C5_15N | 15 | 256 | 0.09 | 0.08 |
| AMP | 15N5 | 15 | 253 | 0.20 | 0.12 |
| biotin | D2 | 15 | 256 | 0.1 | 0.16 |
| betaine | D11 | 11 | 254 | 0.66 | 0.14 |
| pantothenate | 13C3_15N | 0 | 248 | - | 0.20 |
| lysine | D4 | 15 | 256 | 0.50 | 0.33 |
| guanosine | D2 | 12 | 232 | 0.53 | 0.34 |
| 4 aminobenzoic acid | D4 | 15 | 215 | 0.19 | 0.49 |
| cysteine | D3 | 9 | 197 | 0.86 | 0.63 |
